# Twice as nice: Boosts in adolescent reinforcement learning from Pavlovian bias and age-related prioritization of reward-motivated incidental memory

**DOI:** 10.1101/2025.07.11.664442

**Authors:** Haley Hegefeld, Juliet Y. Davidow

**Author notes:** Corresponding author: Juliet Davidow, 360 Huntington Ave, 125 Nightingale Hall, Boston, MA 02115.

## Abstract

Adolescence is a period marked by profound changes in both capacities for learning and the motivational drives that guide behavior. Motivated learning, including the ability to associate cues with actions that lead to positive or negative outcomes, is a fundamental component of adaptive behavior and is essential for survival. Equally important is the encoding of events during learning, which may be influenced by the valence of outcomes. Given the substantial neurocognitive changes in motivated learning and memory that occur from childhood to adulthood, adolescence provides a unique window to investigate mechanisms of these adaptive behaviors. Yet, we know surprisingly little about the development of these behaviors, with sparse extant research fraught with inconsistent findings. In this study, we examined motivated learning and incidental memory using a validated affective learning task in a sample of 174 participants aged 8 to 25 years. The task orthogonalized action and outcome valence and included incidental encoding of trial-unique images presented during feedback, followed by a delayed memory test. We show that adolescents outperform both children and adults in learning by leveraging Pavlovian response biases. In contrast, children exhibit enhanced memory for stimuli associated with positive outcomes compared to adolescents and adults. These findings point to distinct developmental advantages: enhanced learning performance in adolescence and enhanced memory for rewarding events in childhood, each potentially adaptive at their respective developmental stages. Together, these findings suggest opportunities to leverage learning and memory in youth for practical applications, such as education and policy setting.

## Introduction

Learning from experience which actions to take, and which to avoid, is fundamental to adaptive behavior. Simple, robust processes at the core of this can be called motivated learning. Motivated learning is a cognitive building-block, present at the earliest stages of human life (e.g., in infancy; Cuevas & Colombo, 2025; Tummeltshammer et al., 2019), throughout the lifespan (e.g., Chowdhury et al., 2013), and across many species, including rodents (e.g., Wassum et al., 2011; Wiltgen et al., 2007) and nonhuman primates (e.g., Arsenault et al., 2014; Tremblay et al., 1998). Because motivated learning is crucial for an organism’s ability to adapt, studying the factors that can shape this learning is key. Among the greatest influences on motivated learning is the updating of behavior in response to positive and negative outcomes. Updating can be facilitated through Pavlovian learning – the binding of outcomes which reliably follow cues – and instrumental learning – the execution of an action in response to a cue to obtain an outcome. Recent theoretical accounts have noted the dynamic maturation of the neurocognitive systems supporting learning during adolescence, which may give rise to unique benefits for learning guided by feedback, such as occurs in Pavlovian and instrumental learning, during this important life stage (Bolenz et al., 2017; Davidow et al., 2018; Nussenbaum & Hartley, 2019; Wilbrecht & Davidow, 2024).

Pavlovian and instrumental learning have a long history in psychology and neuroscience (Rescorla & Solomon, 1967), are complementary for learning stimulus-outcome and response-outcome associations, respectively, and have well-characterized neural substrates (Averbeck & O’Doherty, 2022; Hart et al., 2014; O’Doherty et al., 2004). In Pavlovian learning, an organism learns to pair a neutral cue with an outcome, such as how a teen learns that a certain “ding” notification tone from their phone means a friend sent a message. Over repeated pairings, the cue (“ding” notification tone) takes on the positive (e.g., “message from a friend”) or negative (e.g., “likely spam”) value of the outcome. These Pavlovian associations can facilitate rapid behavioral responses that depend on the valence of the cue (Guitart-Masip et al., 2014). Cues paired with positive outcomes tend to facilitate approach behaviors while those paired with negative outcomes promote inhibition and withdrawal behaviors, even though the receipt of the outcome is not contingent on the organism’s action (Bouton & Bolles, 1980; Hershberger, 1986; Huys et al., 2011). In this way, the teen will easily learn to look at their phone when they hear their friend’s tone and to *not* look when they hear their parent’s tone after curfew to avoid punishment. These action biases, to approach to gain positive outcomes and to inhibit to avoid negative outcomes, are collectively referred to as “Pavlovian bias.” Pavlovian bias can hinder instrumental learning of incongruent actions, such as learning to approach to avoid negative outcomes. It may be harder for the teen to learn to respond to, rather than to ignore, their parent to avoid a negative outcome for staying out past curfew. Pavlovian influence can bias instrumental learning towards actions that are valence-congruent (i.e., ignoring a parent’s text), even when valence-incongruent actions are advantageous (i.e., responding to the text; Dayan et al., 2006).

In adulthood, Pavlovian bias can boost instrumental learning when congruent, and hinder learning when in conflict (Cavanagh et al., 2013; Guitart-Masip et al., 2012; Huys et al., 2011). We may expect Pavlovian and instrumental learning systems to interact differently as children mature through adolescence and into adulthood (Wilbrecht & Davidow, 2024), even when both systems develop early (Blass et al., 1984; Raab & Hartley, 2018). Adolescents show increased sensitivity to positive outcomes compared to people of other ages (Braams et al., 2015; Galván, 2010), which may impact how adolescents compute value. Heightened sensitivity to positive outcomes can be reflected in behavior, such as greater speeding of responses (Fosco et al., 2022; Galván et al., 2006; Störmer et al., 2014) and physical effort exertion (Rodman et al., 2021), and neurally in greater elicited activation of involved brain regions (Braams et al., 2015; Davidow et al., 2016; Peters & Crone, 2017; Schreuders et al., 2018; Van Leijenhorst et al., 2010) and their connectivity (Davidow et al., 2016; Somerville et al., 2011). Further, youths demonstrate increased impulsivity relative to adults (Doremus-Fitzwater et al., 2012; Harden & Tucker-Drob, 2011; Steinberg et al., 2008), which may alter the relative prepotency of approach vs. withdrawal behaviors. Developmental investigations of Pavlovian influence may elucidate how these hallmark characteristics of cognitive development impact youths’ learning relative to adults’.

Thus far, developmental studies of Pavlovian bias offer conflicting evidence of whether the bias affects learning differently across various ages (Betts et al., 2020; Moutoussis et al., 2018; Raab & Hartley, 2020). An early, large study (*N* = 817) of 14-24-year-olds did not find evidence that Pavlovian influence differs across this age range (Moutoussis et al., 2018). Yet, in a sample of 8-25-year-olds (*N* = 61), adolescents demonstrated less biased learning than children and adults, which led to better performance by adolescents overall (Raab & Hartley, 2020). This result suggested that instrumental learning is less influenced by Pavlovian bias in adolescence than at other ages, allowing for greater flexibility in learning. However, a study of 7-80-year-olds (*N* = 247) found that the influence of Pavlovian bias on instrumental learning gradually increased over the lifespan (Betts et al., 2020). Thus, previous work has found that Pavlovian bias: 1) does not differ across development, 2) attenuates just in adolescence, or 3) increases across the lifespan. These mixed findings warrant further investigation of how Pavlovian bias impacts learning across development. Further, because differences in Pavlovian-instrumental interactions have been identified in several psychiatric disorders (Huys, Gölzer, et al., 2016; Millner et al., 2019; Mkrtchian et al., 2017) and adolescence is a time of peak emergence of these disorders (Lee et al., 2014; Uhlhaas et al., 2023), computational approaches to understanding Pavlovian-instrumental interactions in both normative youth development and in psychiatric populations are critical to advance therapeutic intervention (Hauser et al., 2019; Huys et al., 2021; Huys, Maia, et al., 2016). To enhance our understanding of the impact of learning, we sought to interrogate how outcomes experienced during learning affect memory for incidental features of feedback episodes, thus far unexplored in this prior research.

Outcome associations not only affect behavioral responses, but they also influence what information is prioritized in memory. People tend to remember stimuli associated with positive or negative (vs. neutral) outcomes (Adcock et al., 2006). Further, people tend to remember positively and negatively surprising events, or when there is a difference between the outcome they expect to receive and what they actually receive, called a prediction error (Calderon et al., 2021; Jang et al., 2019; Pupillo et al., 2023; Rouhani & Niv, 2021). The association between prediction error and memory may be particularly strong during development, because adolescents are more attuned to positive outcomes, which could enhance memory encoding of stimuli associated with unexpectedly positive outcomes during this period of life (Braams et al., 2015; Galván, 2010). One previous study of adolescents and adults supports the hypothesis that adolescents show a stronger bias in their memory towards stimuli associated with unexpectedly positive (vs. negative) outcomes than adults do, which may be due in part to maturational differences in the neural substrates supporting learning (Davidow et al., 2016). The current study seeks to test this hypothesis in a larger sample with a wider developmental age range extending from middle childhood to young adulthood.

To address how Pavlovian bias impacts instrumental learning during development, we had 174 participants, 8–25 years of age (see Methods and Fig. S1 for detailed demographics), complete a probabilistic reinforcement learning task that orthogonalizes action and outcome valence, the “Affective Go/No-Go Task” (Guitart-Masip et al., 2012; Figs. 1A and 1B). Participants learned by trial and error which of four cues signaled that the participant should: 1) press the “space” key to win money (i.e., “Go to Win”), 2) press to avoid losing money (i.e., “Go to Avoid Loss”), 3) not press to win money (i.e., “No-Go to Win”), and 4) not press to avoid losing money (“No-Go to Avoid Loss”). Pavlovian bias and instrumental learning are congruent in the Go to Win and No-Go to Avoid Loss conditions, and incongruent in the No-Go to Win and Go to Avoid Loss conditions (Fig. 1C). If Pavlovian bias impacts instrumental learning, participants should perform better in the conditions in which Pavlovian and instrumental learning are congruent than in those in which they are incongruent. To address how positive and negative outcomes affect incidental memory encoding during development, we displayed a trial-unique image of an object during outcome presentation in the learning task, and surprised participants with a memory test later in the study session.

**Figure 1.**
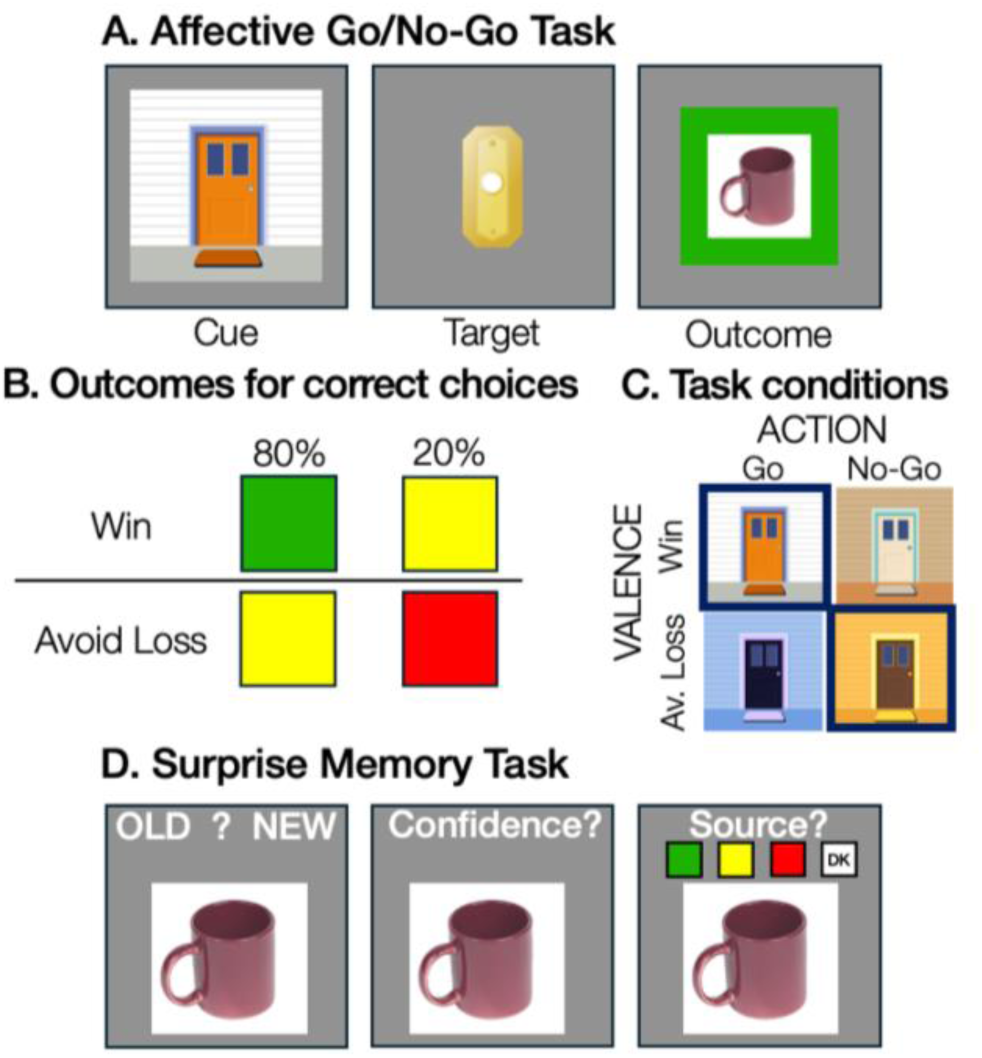
(A) In the Affective Go/No-Go Task, participants first see one of four doors (the cue), after which they choose whether to ring the doorbell (the target), and then receive feedback according to their choice (the outcome). (B) For doors signaling the potential to win money, 80% of the time participants will win $0.25 for making the correct choice (depicted as a green frame around a trial-unique image of an everyday object) and win $0 for making the incorrect choice (yellow frame). For doors signaling the potential to lose money, 80% of the time participants will lose $0 for making the correct choice (the yellow frame) and will lose $0.25 for making the incorrect choice (red frame). The feedback is probabilistic, such that the outcome is reversed on 20% of trials. (C) Action and valence are orthogonalized across the four doors. Thus, each door signals the correct action, in addition to the potential to win/lose money. Pavlovian-congruent conditions are outlined by a navy box, while Pavlovian-incongruent conditions are not. (D) Participants complete a surprise memory task of the trial-unique images of objects presented during the outcome phase of the learning task interleaved with novel object images. Participants first identify whether an image is “Old” or “New”, then they rate their confidence and report their source memory for the associated outcome.

We hypothesized that instrumental learning in adolescence would exceed performance in younger and older participants, based on prior work demonstrating a boost in instrumental learning in adolescence compared to other ages (Davidow et al., 2016; Raab & Hartley, 2020), which may be due to adolescents’ higher sensitivity to positive outcomes during motivated learning than other ages’ (Casey et al., 2025). We also sought to elucidate the influence of Pavlovian bias on different conditions of instrumental learning given mixed findings in prior developmental literature. Because unexpectedly positive outcomes have been shown to enhance encoding of stimuli incidentally associated with these outcomes in adolescents (Davidow et al., 2016), we hypothesized that younger participants would show a stronger prioritization of images associated with unexpectedly positive outcomes in memory than adult participants would. Overall, this study sought to examine potential cognitive benefits that adolescents experience in motivated learning and incidental memory during this unique phase of neurocognitive development.

## Results

### Pavlovian-congruency boosts instrumental learning in adolescence

We examined whether adolescents demonstrated better instrumental learning than children and adults. To address this central question, we employed mixed-effects logistic regression (Brooks et al., 2017). We compared nested models with task-related and age-related predictors. For age, we tested whether instrumental learning showed an adolescent-specific effect, which would support our hypothesis, a gradual developmental effect, or a non-developmental effect by comparing quadratic, linear, and null (i.e., not including age) coefficients, respectively. The best-fitting regression model (ΔBIC for next-best regression model = 19, see “FE Quadratic Age - Parsimonious”, Table S2) included fixed-effects for trial number, action and valence conditions, and linear and quadratic age, with random intercepts for participants (Table 1, Fig. S2).

**Table 1.** Results from the Best-Fitting Generalized Mixed-Effects Model of Learning Accuracy.

| Fixed Effects |  |  |  |  |
| --- | --- | --- | --- | --- |
|  | Odds Ratio | 95% CI | Z-value | p-value |
| Intercept | 1.31 | 1.18-1.45 |  |  |
| Trial | 1.38 | 1.31-1.45 | 13.21 | < .001 |
| Action (Ref: No-Go) | 1.44 | 1.35-1.54 | 10.62 | < .001 |
| Valence (Ref: Avoid Loss) | 0.55 | 0.52-0.59 | -17.49 | < .001 |
| Age | 1.21 | 1.10-1.34 | 3.81 | < .001 |
| Age<sup>2</sup> | 0.78 | 0.70-0.86 | -4.92 | < .001 |
| Trial x Action | 0.72 | 0.68-0.77 | -9.40 | < .001 |
| Trial x Valence | 0.91 | 0.85-0.97 | -2.76 | .006 |
| Action x Valence | 4.21 | 3.81-4.65 | 28.10 | < .001 |
| Trial x Age | 1.07 | 1.03-1.11 | 3.94 | < .001 |
| Trial x Age<sup>2</sup> | 0.99 | 0.96-1.03 | -0.34 | .733 |
| Action x Age | 0.83 | 0.77-0.88 | -5.64 | < .001 |
| Action x Age<sup>2</sup> | 1.27 | 1.18-1.35 | 6.89 | < .001 |
| Valence x Age | 0.90 | 0.84-0.96 | -3.22 | .001 |
| Valence x Age<sup>2</sup> | 1.17 | 1.09-1.25 | 4.61 | < .001 |
| Trial x Action x Valence | 1.37 | 1.24-1.52 | 6.21 | < .001 |
| Trial x Action x Age | 0.94 | 0.89-0.98 | -2.69 | .007 |
| Trial x Action x Age<sup>2</sup> | 0.91 | 0.87-0.96 | -3.56 | < .001 |
| Action x Valence x Age | 1.18 | 1.07-1.30 | 3.28 | .001 |
| Action x Valence x Age<sup>2</sup> | 0.66 | 0.60-0.73 | -8.28 | < .001 |
| Marginal pseudo-*R*<sup>2</sup> (Fixed Effects): 12.7% |  |  |  |  |
| Conditional pseudo-*R*<sup>2</sup> (Fixed and Random Effects): 21.1% |  |  |  |  |
| Random Effects |  |  |  |  |
|  | Variance | Standard Deviation |  |  |
| Subject (Intercept) | 0.35 | 0.59 |  |  |
|  | Model Equation: Accuracy ~ Trial + Action + Valence + poly(Age,2) + Trial*Action + Trial*Valence + Action*Valence + Trial*poly(Age,2) + Action*poly(Age,2) + Valence*poly(Age,2) + Trial*Action*Valence + Trial*Action*poly(Age,2) + Action*Valence*poly(Age,2) + (1\|Subject) |  |  |  |

The effect of Pavlovian bias on instrumental learning is characterized by an interaction of Action*Valence (Fig. 2A), which was significant in the best-fitting regression model (*p* < .001) and was qualified by a significant interaction of Action*Valence*Quadratic Age (*p* < .001), suggesting adolescent-specific performance effects for some action-valence conditions. The best-fitting model also included a significant interaction for Action*Valence*Trial Number (*p* < .001), indicating that there may be differences in learning curves for some conditions, unrelated to age (see Figs. S3-S5 for interaction plots). To determine whether the findings from these complex interactions supported our hypothesis that learning here is enhanced in adolescents, we conducted follow-up beta-binomial regression models that assessed whether learning accuracy summarized across all trials and summarized within each action-valence condition were predicted by participants’ ages.

**Figure 2.**
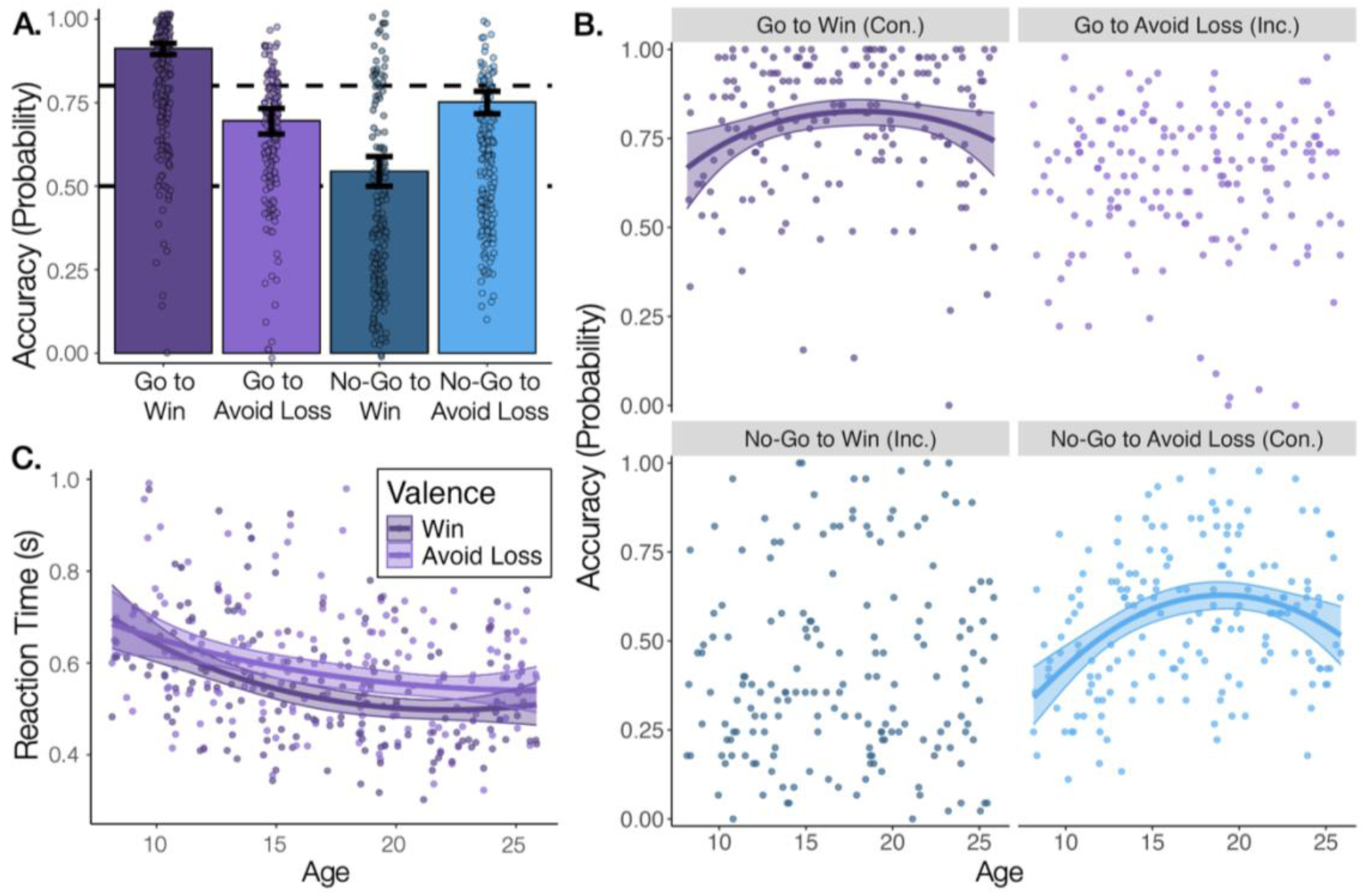
(A) Pavlovian influence on instrumental learning is evident at the group level. Participants perform better in the Pavlovian-congruent than Pavlovian-incongruent condition within each action (i.e., Go to Win > Go to Avoid Loss; No-Go to Avoid Loss > No-Go to Win, see Fig. S3 for paired t-tests). Bars display model predictions for the last trial of the task at the mean age of participants (Fig. S5 shows plots at different trials and ages). Error bars show 95% confidence intervals, and points show each participant’s accuracy within the trial types averaged across the entire task. Dashed lines at 50% and 80% represent chance and probability-matching performance, respectively. (B) There is a peak in performance for adolescents in the Pavlovian-congruent conditions (“Con.”), whereas participants of all ages perform similarly in the Pavlovian-incongruent conditions (“Inc.”). (C) Adolescents demonstrate the greatest relative speeding to the Pavlovian-congruent Go to Win vs. Pavlovian-incongruent Go to Avoid Loss conditions, shown in the plot by the difference between the two fit lines. In (B) and (C) the lines show model predictions across age, shaded regions show 95% confidence intervals, and points show the participants’ raw data averaged across the task within relevant conditions. Fit lines are shown for statistically significant effects.

We found that overall learning accuracy summarized across all trials was best predicted by a quadratic age effect (Linear vs. Null: *χ^2^(1)* = 3.54, *p* = .06; Quadratic vs. Linear: *χ^2^(1)* = 10.58, *p* = .001). Both linear and quadratic age were significant predictors in this model (Linear Age: Odds Ratio (*OR*) = 1.08, 95% CI = [1.00, 1.16], *z* = 1.95, *p* = .05; Quadratic Age: *OR* = 0.88, 95% CI = [0.81, 0.95], *z* = −3.31, *p* < .001), supporting our hypothesis that adolescents would show enhanced instrumental learning compared to younger and older participants.

This well-used paradigm consistently finds variability in learning that reflects individual differences in motivational drives. Broadly these differences in motivation consist of sensitivity to the potential to gain rewards compared to avoiding losses, and to the congruency between the Pavlovian and instrumental states. We sought to understand how the adolescent peak in overall learning we observed was supported by these motivational drives. As such, we examined how age predicted learning accuracy within each action-valence condition, which orthogonalizes the influence of gains vs. losses and Pavlovian-congruency (Fig. 2B). For the Go to Win condition, which is Pavlovian-congruent, accuracy was best predicted by a quadratic age effect (Linear vs. Null: *χ^2^(1)* = 1.24, *p* = .27; Quadratic vs. Linear: *χ^2^(1)* = 5.79, *p* = .02). Quadratic age was a significant predictor in this model (*OR* = 0.82, 95% CI = [0.70, 0.96], *z* = −2.46, *p* = .01), but linear age was not (*OR* = 1.09, 95% CI = [0.94, 1.27], *z* = 1.12, *p* = .26). For the No-Go to Avoid Loss condition, which is also Pavlovian-congruent, the quadratic age model fit the data significantly better than the linear age model, which outperformed the null (Linear vs. Null: *χ^2^(1)* = 9.37, *p* = .002, Quadratic vs. Linear: *χ^2^(1)* = 15.44, *p* < .001). Both linear (*OR* = 1.20, 95% CI = [1.07, 1.34], *z* = 3.23, *p* = .001) and quadratic age (*OR* = 0.80, 95% CI = [0.71, 0.89], *z* = −3.99, *p* < .001) were significant predictors in this model. Neither the linear nor the quadratic age models performed better than the null model for the Pavlovian-incongruent conditions, Go to Avoid Loss and No-Go to Win (*ps* > .20). This suggests a benefit for learning in adolescents from Pavlovian-congruency even in loss avoidance contexts. Reaction time also demonstrated a quadratic age pattern (Quadratic vs. Linear Age Model: *χ^2^*(2) = 10.8, *p* < .01), such that adolescents showed the greatest relative speeding of their responses to the Pavlovian-congruent Go to Win vs. the Pavlovian-incongruent Go to Avoid Loss stimuli, compared to children and adults (Fig. 2C). Overall, these results suggest that adolescents perform better than people of other ages in the Pavlovian-congruent conditions, and people of all ages perform similarly in the Pavlovian-incongruent conditions.

### Mechanisms of adolescent boost in learning

Given that we saw adolescent boosts in learning in Pavlovian-congruent conditions, we investigated Pavlovian bias through behavioral and reinforcement learning approaches to further support this interpretation. We computed a behavioral metric of Pavlovian bias (Cavanagh et al., 2013; Raab & Hartley, 2020), which reflects a participant’s tendency to “Go” for cues with the potential to win money and “No-Go” for cues with the potential to lose money, the two Pavlovian-congruent action biases. Complementing the results of the regression, the linear age model of Pavlovian bias did not fit significantly better than the null model (*F*(1,172) = 0.45, *p* = .50), but the quadratic age model fit better than the linear age model (*F*(1,171) = 4.90, *p* < .05; Fig. S7). This suggests that adolescents’ learning may be more influenced by Pavlovian bias than people of other ages’.

To formalize putative cognitive mechanisms underlying the task behavior, such as Pavlovian bias, we fit a set of reinforcement learning (RL) models to the data using an expectation maximization approach (Huys, 2018; Huys et al., 2011; Sutton & Barto, 1998). This approach fits models hierarchically, which allows the group-level and individual-level parameter distributions to iteratively constrain each other instead of setting priors for parameters. This approach can be considered conservative for a search of individual differences, such as age-related differences, as individual parameter solutions are constrained towards the group mean. We selected a set of models to aid in our interpretation of the age-related differences in behavioral findings reported above, and other models that would permit us to make direct comparisons with prior literature (see Table S4 for model list). Performance here was best captured by a RL model (“Valenced Sensitivity + Pavlovian Bias” model, Table S4) that included a single learning rate, separate parameters for gain and loss sensitivities, “go” bias, Pavlovian bias, and a lapse rate. This model fit the younger participants slightly better than the older participants, as suggested by a weak, positive correlation between a model comparison metric estimated for each participant (i.e., BIC; Schwarz, 1978) and their age (*r* = .15, 95% CI = [.001 .29], *t*(172) = 1.99, *p* = .048; Fig. S10). Though there was this age-related trend in model fit, given our hypotheses we considered age associations with model parameters to determine whether certain latent cognitive mechanisms supported the adolescent peak in performance for Pavlovian-congruent conditions. The only parameter that was associated with age was loss sensitivity, which was slightly higher in the older participants than in the younger (*r* = .20, 95% CI = [.05, .34], *t*(172) = 2.66, *p* < .01; see Fig. S10 for all parameter associations with age).

Notably, though behavioral results suggested that adolescents exploit a higher Pavlovian bias during instrumental learning, the Pavlovian bias parameter from the best-fitting RL model did not suggest an adolescent enhancement. Finally, we sought to understand whether incidental features that co-occur with positive outcomes lead to enhanced memory in children, adolescents, and adults.

### Reward enhanced memory in youth

To investigate whether incidental memory encoding is enhanced by positive outcomes across ages, we computed *d’* as a measure of overall accuracy that accounts for the participant’s tendency to report that they saw an image (Stanislaw & Todorov, 1999). We also tested for a reward memory bias, by examining age-related differences in *d’* for images shown when the participants could gain a reward and did gain, and when they could gain a reward but did not during the learning task. For overall *d’*, the linear age model fit significantly better than the null model (Linear vs. Null: *F*(1,160) = 24.22, *p* < .001; Age: *β* = −0.36, 95% CI = [−0.51, −0.22], *t*(160) = −4.90, *p* < .001). This indicated that increasing age is associated with relatively poorer performance on the memory task (Fig. 3A). Importantly, we found an interaction of reward and linear age on *d’*, such that younger participants remembered images paired with gaining a reward better than not gaining a reward, while older participants showed the opposite (Reward*Age: *β* = −0.10, 95% CI = [−0.15, −0.04], *z* = −3.44, *p* < .001; Figs. 3B and S11). This showed that younger participants’ memory of images associated with rewards, in particular, is better than adults’. Given that we found age-related differences in both learning and memory performance, we also assessed whether learning and memory accuracy had a trade-off, which may indicate that learning from feedback and incidental memory encoding competed for resources, and found a weak, negative correlation (*r* = -.17, 95% CI = [−.32, -.02], *t*(160) = −2.20, *p* = .03; Fig. S12). This suggests that the enhanced memory performance by younger participants here could be explained by learning-memory trade-off.

**Figure 3.**
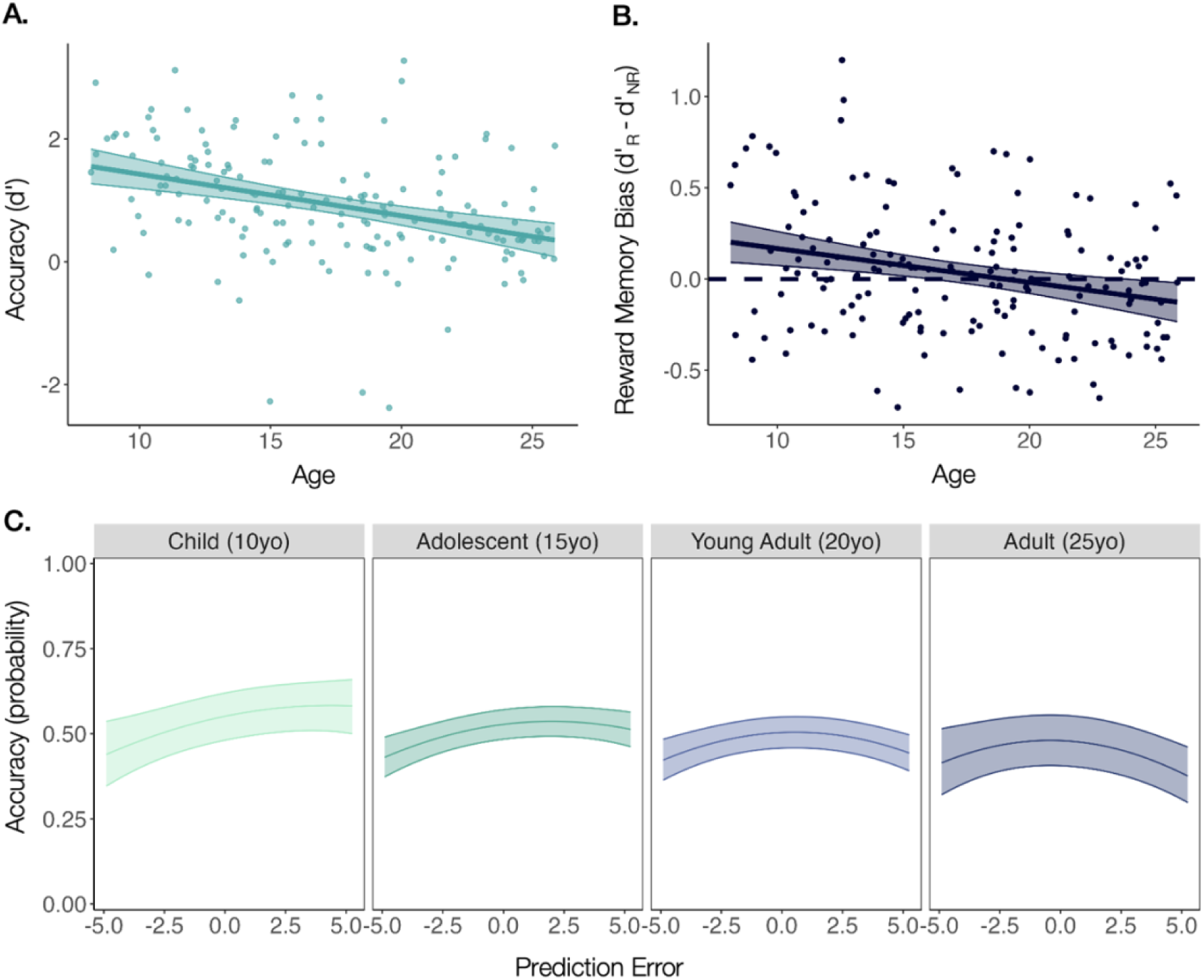
(A) Memory accuracy (*d’*) negatively correlates with age. Shaded regions show 95% confidence intervals, and points show the raw data. (B) We visualized the interaction of Reward-by-Age on d’ by plotting a difference score here, showing that reward memory bias is higher in the younger participants than the older. See Fig. S11 for the effects plot of the interaction. R = Reward, NR = No Reward. (C) The Prediction Error-by-Age interaction shows that children have positive prediction error-related memory enhancements relative to adolescents and adults. The lines represent model predictions at each age (10, 15, 20, and 25-years-old), and shaded regions show 95% confidence intervals.

Prediction errors in reinforcement learning are the result of expectancy violations, and can capture attentional resources in service of updating one’s mental model. Thus, prediction errors may increase the likelihood that stimuli presented at the same time are attended to and encoded in incidental memory. Indeed, prior work has demonstrated that prediction error can elicit both signed (i.e., better memory for positively surprising events) and unsigned (i.e., better memory for surprising events regardless of valence) effects on memory (Ergo et al., 2020; Pupillo et al., 2023). We expected adolescents to show a stronger signed effect of prediction error on memory, such that they remember images associated with positively (vs. negatively) surprising outcomes better than people of other ages, based on prior work showing a similar age-related difference (Davidow et al., 2016).

To test our central question about how prediction error and memory are integrated at different ages, we extracted trial-by-trial prediction errors from the RL model. We submitted the prediction errors and age to mixed-effects logistic regression models predicting trial-by-trial accuracy for previously seen images. We tested models that included a predictor for linear prediction error (to account for signed prediction errors) and quadratic prediction error (to account for unsigned prediction errors). The best-fitting regression model included fixed-effects for linear and quadratic prediction error, and linear age, with random intercepts for participants (Table S5). Model results are presented in Table 2. The significant Prediction Error*Age interaction (*p* < .001) suggests that there are age-related differences in the association between prediction error and memory accuracy. To understand these age-related differences, we visualized the association between prediction error and memory accuracy at representative ages (Fig. 3C). Children’s memory accuracy improved as prediction errors became increasingly positive (i.e., a signed effect), suggesting a prioritization of images appearing with outcomes that were more rewarding than expected. At older ages, this association was attenuated. Together, these results complement prior work showing that increased sensitivity to gaining rewards may have benefits for memory especially during development.

**Table 2.**
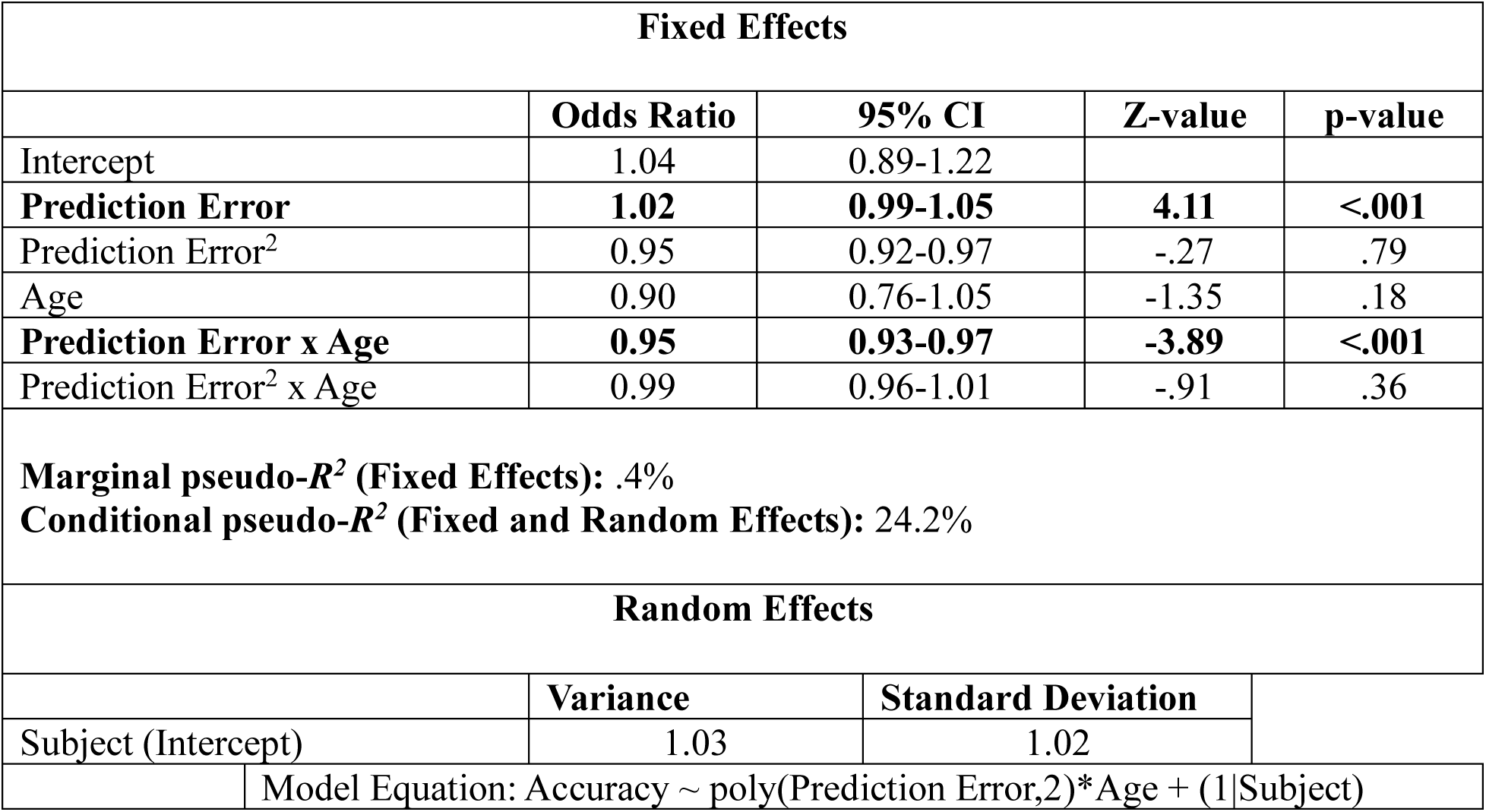
Results from the Best-Fitting Generalized Mixed-Effects Model of Memory Accuracy.

| Fixed Effects |  |  |  |  |
| --- | --- | --- | --- | --- |
|  | Odds Ratio | 95% CI | Z-value | p-value |
| Intercept | 1.04 | 0.89-1.22 |  |  |
| Prediction Error | 1.02 | 0.99-1.05 | 4.11 | <.001 |
| Prediction Error<sup>2</sup> | 0.95 | 0.92-0.97 | -.27 | .79 |
| Age | 0.90 | 0.76-1.05 | -1.35 | .18 |
| Prediction Error x Age | 0.95 | 0.93-0.97 | -3.89 | <.001 |
| Prediction Error<sup>2</sup> x Age | 0.99 | 0.96-1.01 | -.91 | .36 |
| Marginal pseudo-*R*<sup>2</sup> (Fixed Effects): .4% |  |  |  |  |
| Conditional pseudo-*R*<sup>2</sup> (Fixed and Random Effects): 24.2% |  |  |  |  |
| Random Effects |  |  |  |  |
|  | Variance | Standard Deviation |  |  |
| Subject (Intercept) | 1.03 | 1.02 |  |  |
| Model Equation: Accuracy ~ poly(Prediction Error,2)*Age + (1\|Subject) |  |  |  |  |

## Discussion

In this study, we investigated how the reflexive Pavlovian responses to approach to gain positive outcomes and to withhold to avoid negative outcomes impact instrumental learning across development. We also sought to understand how reward learning constrains what is prioritized in memory at different ages. We found that Pavlovian influence is evident across development, and that adolescents may be particularly adept at maximizing outcomes in such contexts. Children remembered incidentally-encoded images better than adolescents and adults, which was driven by an increased prioritization of images associated with unexpected rewards.

Our findings generally supported our hypothesis that adolescents would show instrumental learning benefits. Using logistic regression, we demonstrated that adolescents performed better than children and adults only in Pavlovian-congruent conditions. This finding was complemented by a behavioral metric of Pavlovian bias that peaked in adolescence. However, the Pavlovian bias parameter estimated by a computational RL model was not associated with age. There are a few possible explanations for this seeming inconsistency. For one, we had a weak association between age and model fit, which could have suggested that different models would have fit different ages better. To be conservative in the number of different fitting approaches and to avoid arbitrary cut-offs to bin ages into groups, we did not explore this. For another, because we fit the RL model hierarchically, we may have skewed parameter solutions towards a group mean that diminished age-related differences in the Pavlovian bias parameter. In either case, multiple behavioral analysis approaches showed that Pavlovian influence was stronger in adolescence than at other ages, which strengthens the interpretation of the adolescent-specific boosts even without complementary RL results.

Better learning in Pavlovian-congruent conditions across ages is consistent with previous developmental studies that show Pavlovian responses to approach to gain positive outcomes and to withhold to avoid negative outcomes influence instrumental learning in childhood, adolescence, and adulthood (Betts et al., 2020; Moutoussis et al., 2018; Raab & Hartley, 2020). Yet, this prior literature is inconsistent in drawing precise conclusions regarding the interaction between Pavlovian and instrumental learning systems for adolescents. Here we outline these inconsistencies and suggest interpretations to reconcile these differing patterns.

The first inconsistency is that adolescents performed better than people of other ages in the current study, which concurs with one previous study (Raab & Hartley, 2020) but conflicts with another that found superior performance by young adults (Betts et al., 2020). Perplexingly, adolescents performed especially well in the Pavlovian-incongruent conditions in the previous study finding adolescent performance boosts (Raab & Hartley, 2020), but in the Pavlovian-congruent conditions in the current study.

A second inconsistency is the best-fitting RL model for participant choice behavior, which also differs by study. Our winning RL model included the same parameters—a learning rate, lapse rate, gain and loss sensitivities, go bias, and Pavlovian bias—as in a previous study (Betts et al., 2020). However, the winning model in another study had only one reinforcement sensitivity but valenced learning rates (Moutoussis et al., 2018; although a valenced sensitivity model also captured choice behavior well), while yet another winning model had only one reinforcement sensitivity and one learning rate (Raab & Hartley, 2020). Models with distinct parameters for gains and losses, whether that be valenced learning rates (Hauser et al., 2015; Moutoussis et al., 2018; Rosenbaum et al., 2022; van den Bos et al., 2012) or valenced outcome sensitivities (Betts et al., 2020; Cavanagh et al., 2013), frequently outperform models that have one parameter for both, suggesting that people may learn differently from positive and negative outcomes (Nussenbaum & Hartley, 2019). Consistent with this view, we found that loss sensitivity, but not gain sensitivity, weakly increased across age. This is perhaps a third inconsistency, as a previous study found that both gain and loss sensitivity are highest in young adults relative to children, adolescents, and older adults (Betts et al., 2020).

A fourth inconsistency is the age trajectory of Pavlovian bias. Our cross-sectional study found that the Pavlovian bias parameter estimated by a computational model did not differ by age. This is consistent with the only longitudinal study, which found no change in Pavlovian bias across age (Moutoussis et al., 2018), but inconsistent with the other cross-sectional studies that found an attenuated Pavlovian bias in adolescence (Raab & Hartley, 2020) or a gradual increase in the bias over the lifespan (Betts et al., 2020). How could we reconcile these inconsistencies?

One potential explanation is that Pavlovian response biases are not rigid reactions, but rather operate like prior beliefs. This explanation is supported by the results of the only longitudinal investigation of Pavlovian bias (Moutoussis et al., 2018). Because the study was longitudinal, the authors were able to disentangle age-related and practice-related effects. The authors found that Pavlovian bias decreases over test sessions, but not over age. This was interpreted as evidence that the bias operates as a prior belief (e.g., “If I have the possibility to gain a reward, the correct decision is to Go.”) that can be overcome with experience, rather than as a disposition that changes across development. It is possible that adolescents in the current study exploited this Pavlovian prior belief when they found that it resulted in advantageous outcomes for some stimuli, resulting in better performance in Pavlovian-congruent conditions. These adolescents may have needed more experience (i.e., trials) to overcome the prior for the Pavlovian-incongruent conditions. Future work could systematically vary the number of trials in the learning task or include task repetitions to examine how more or less experience impacts learning at different ages.

Another explanation is that instrumental learning is a flexible system that is highly sensitive to context. Theoretical expositions have suggested that this may account for inconsistencies observed in the associations between age and common RL parameters (Nussenbaum & Hartley, 2019). The authors suggested that the true developmental difference in RL may be an age-related enhancement in an individual’s ability to adjust their learning parameters to a specific environment, rather than a simple increase or decrease in a particular parameter. Although this hypothesis does not fully account for why adolescents performed better than adults in the current study, it highlights that RL parameters are influenced by one’s context and are far from stable, trait-like qualities. A recent study tested this hypothesis by comparing RL parameters from three tasks within-subjects. The study found low to moderate generalizability in the RL parameters across tasks (Eckstein et al., 2022). As the task versions used in previous studies and the current study all differ in length, cover story, and reward magnitudes, we may expect that diverging results would emerge as participants of different ages adjust their learning to the current environment. Given that the study about parameter generalizability suggests that parameters may differ in how much they are adjusted to a specific environment, future research should take a similar approach to understanding whether Pavlovian bias is context-dependent, as has been suggested by studies manipulating the controllability of the environment (Dorfman & Gershman, 2019; Raab et al., 2024).

Turning to the memory results, children remembered incidentally-encoded images better than adolescents and adults and privileged memories for images associated with better-than-expected outcomes. Though an age-related decrease in memory sensitivity was not hypothesized, another developmental study that tested memory for incidentally-encoded images also observed a negative association between age and memory accuracy (Rosenbaum et al., 2022). In the current study, children’s superior memory was more pronounced for images associated with positive vs. negative prediction errors, indicating an effect of signed prediction error on memory encoding in childhood. This is consistent with a burgeoning literature that suggests prediction error may tag events that ought to be remembered, as these events may be particularly informative (Calderon et al., 2021; Jang et al., 2019; Pupillo et al., 2023; Rouhani & Niv, 2021). However, most of these studies have focused on adults, who did not show strong prediction error-related memory enhancements in the current study. One study of adolescents and adults also found an effect of signed prediction error on memory in adolescence, such that adolescents had better memory for incidentally-encoded images paired with positive prediction errors than adults (Davidow et al., 2016), but did not investigate children younger than 13-years-old. Our findings, complemented by prior research (Davidow et al., 2016; Rosenbaum et al., 2022), preliminarily suggest that prediction error exerts a signed effect on memory during development, promoting memory for better-than-expected outcomes.

There are several potential explanations for the age-related pattern observed here, such that children show stronger prediction error-related memory enhancements than adolescents and adults. One possibility is that reinforcement learning and memory competed for cognitive resources, as suggested by the trade-off in learning and memory accuracy found here and by a previous study of adults that showed increased memory encoding interfered with learning (Wimmer et al., 2014). However, this does not explain why adults’ had relatively lower performance on both the learning and memory tasks. Another possibility is that individual differences qualify this age-related association. One developmental study revealed that the extent to which an individual prioritizes better-than-expected outcomes (i.e., the difference in their learning rates for positive and negative outcomes) is associated with their memory for images paired with those outcomes (Rosenbaum et al., 2022). Although a two-learning rate model did not fit the observed data best here, it is possible that children weighted positive prediction errors more than adolescents and adults, resulting in better memory for those events. More studies investigating the interactions between learning and memory systems from childhood to adulthood are needed to draw conclusions.

Future research examining how the neural substrates of Pavlovian-instrumental learning interactions, as well as of incidental memory encoding, change across development may help to explain the various age-related differences found in this study. Prior work has shown that Pavlovian and instrumental learning are supported by distinct cortico-striatal circuits, both of which are predominantly associated with dopaminergic signaling (Averbeck & O’Doherty, 2022; O’Doherty et al., 2004; Yin et al., 2005). A previous fMRI study of adults who completed the Affective Go/No-Go Task found that participants who successfully learned had more activation in the substantia nigra/ventral tegmental areas during anticipation of “Go” trials, regions that track prediction errors through firing rates of dopamine neurons and project to the striatum (Schultz et al., 1997), and in the inferior frontal gyrus during anticipation of “No-Go” trials, a region which supports response inhibition (Aron et al., 2014). Further, two EEG studies of adults demonstrated that midfrontal theta power supported the ability to overcome Pavlovian bias trial-by-trial during this task (Cavanagh et al., 2013; Gershman et al., 2021). Given that cortico-subcortical and cortico-cortical connections continue to develop throughout adolescence and into young adulthood (Casey et al., 2019; Davidow et al., 2019; van Duijvenvoorde et al., 2019), it is possible that the maturational stage of adolescents’ cortico-striatal circuitry bolstered the tendency to exploit Pavlovian bias at these ages. It is also possible that the incidental memory encoding during feedback presentation recruited hippocampal and medial temporal lobe systems to a greater extent in the younger participants, as found by Davidow and colleagues (2016), supporting the learning and memory benefits of the adolescents and children, respectively. Future neuroimaging studies may reconcile the findings discussed here by uncovering how neural circuitry is recruited to support learning and memory across development. Longitudinal investigations will be especially helpful for this effort, as much of the prior work, including this study, is limited in interpreting developmental trajectories by using cross-sectional samples.

In conclusion, we examined how Pavlovian influence constrains instrumental learning across development, as well as how learning influences later memory. Adolescents demonstrated peak learning performance relative to children and adults, which was driven by better accuracy in the conditions facilitated by Pavlovian bias. Our results are consistent with and differ from results of previous developmental studies of Pavlovian bias in several ways. We identify avenues for future research to better understand these inconsistencies, the most pressing of which may be to understand how contextual factors influence Pavlovian bias and its developmental trajectory. Children had stronger positive prediction error-related memory enhancements than adolescents and adults. Interactions between Pavlovian and instrumental learning systems, and between instrumental learning and episodic memory, may reveal how the brain scaffolds learning and memory at different stages of neurodevelopment, critical for furthering basic and practical applications in many domains. Our results join with a growing body of research revealing unique cognitive advantages during development (Casey et al., 2025), reinforcing that research must continue to move away from models that promote a “broken brain” framing. Moving forward, we must work to leverage these advantages for practical applications, such as in classroom or clinical interventions.

## Methods

### Participants

Participants were recruited for this online study from across the United States using online advertisements. The eligible age range was 8–25 years old, and participants were screened to exclude for mood or anxiety disorder diagnoses, learning disabilities, current use of psychoactive medications, color-blindness, low proficiency with English, and lack of technology necessary to complete an online study from home. 174 eligible participants completed the RL task (*n* = 87 female, 84 male, 3 non-binary, mean age = 17.09 years, SD = 5 years), and a subset of 162 eligible participants also provided usable memory task data (exclusions described in the “Data Processing” section, *n* = 79 female, 80 male, 3 non-binary, mean age = 17.18 years, SD = 4.95 years). Of the 174 participants, 90 identified as White (51.7%), 55 as Asian (31.6%), 18 as Black/African American (10.3%), 6 as multiracial (3.45%), 2 as other (1.15%), and 3 individuals did not disclose race. 12 participants identified as Hispanic or Latino (6.9%). All study procedures were approved by the university ethics committee. Participants were paid $12/hr for their time, with additional earnings based on performance in the reinforcement learning task.

### Procedure

All study procedures were conducted during a HIPAA-compliant Zoom video call with an experimenter. After providing informed consent or assent, participants first completed the RL task. Participants completed the surprise memory task approximately 30 minutes after finishing the learning task. In between the tasks, the experimenter administered the Wechsler Abbreviated Scale of Intelligence (WASI-II; Wechsler, 2011) Vocabulary and Matrix Reasoning subtests, and participants answered a battery of self-report questionnaires. After completing the memory task, adult participants finished remaining questionnaires as needed, and all participants were debriefed and compensated. To reduce fatigue, minor participants could complete questionnaires on their own time in the weeks following the study for additional payment.

#### RL Task Details

The Affective Go/No-Go Task (Guitart-Masip et al., 2012) requires participants to learn whether to respond or withhold a response, from outcomes where they gain or avoid losing $0.25. We implemented a cover story where participants were instructed to collect as many donations for a charity as possible, which required participants to learn whether they should ring or not ring a doorbell (i.e., pressing the space key or withholding the press) for each door. Mapping of four unique door images to the trial types (i.e., Go to Win, Go to Avoid Loss, No-Go to Win, and No-Go to Avoid Loss) was counterbalanced across participants. Participants were instructed that they would also earn money for themselves by collecting donations, which they received as actual earnings at the end of the study session.

On each trial, participants saw the cue (i.e., one of four doors, 1.5s), followed by a fixation cross (jittered .4-1.2s), and then the target (i.e., the doorbell, 1.5s; Fig. 1A). When participants saw the target, they chose whether to press the space key (“Go” response) or not (“No-Go” response). Participants then saw a fixation cross (jittered .2-.6s), followed by the probabilistic outcome (2s). A trial-unique image of an everyday object was presented with the outcome, which was used in the subsequent memory task. During the intertrial interval, a fixation cross appeared on the screen (.7s).

Participants completed 45 trials of each type, for a total of 180 trials. The trial order was pseudorandomized, such that there were 15 trials of each type in each of three blocks.

Participants had a self-paced break between each block. Prior to beginning the task, participants completed eight instructed practice trials where they made a “Go” and “No-Go” response to four doors (which were grayscale to distinguish from doors used during the task) to experience winning, not winning nor losing, and losing money depending on their choice, followed by eight uninstructed practice trials. Participants were instructed about the probabilistic nature of the feedback, although they were not told the exact probability.

#### Surprise Memory Task Details

Participants completed a surprise memory test of all the trial-unique images presented in the learning task and an additional 90 novel images, for a total of 270 images (Fig. 1D). On each trial, participants first indicated whether they remembered seeing an image in the learning task (responses: *Old, New*). Then, they reported their confidence in their decision (responses: *Guessing, Pretty Sure, Very Sure, Completely Certain*). Finally, they chose the associated outcome on that trial if they selected “Old” or had to choose “IDK” if they selected “New.” All decisions were self-paced. Trial order and assignment of images to the “Old” and “New” conditions were randomized across participants. Both experimental tasks were programmed in PsychoPy (v. 2020.2.0) to be run online through the Pavlovia server (Peirce et al., 2019).

### Data Processing

Data processing and analyses were conducted in R (v. 4.3.1; R Core Team, 2023), and deidentified study data and analysis scripts are available on the Open Science Framework (*link to be added upon publication*). Minimal processing of the task data was completed using the tidyverse package (i.e., removing practice trials) (Wickham et al., 2019). Each learning trial was counted as “correct” if the participant made the optimal action associated with the door regardless of the outcome they saw. In other words, even if the participant saw the probabilistically rare outcome of a yellow screen after making the correct “Go” response for a “Go to Win” door, we tallied this as a “correct” response. To measure memory performance, we counted trials as “correct” when the participant accurately identified a signal image as being “Old” (a “hit”) or when they accurately identified a novel image as being “New” (a “correct rejection”). An incorrect trial was either a “miss” (i.e., identifying a signal image as “New”) or a “false alarm” (i.e., identifying a novel image as “Old”).

We censored data in the memory task by balancing the necessity to: 1) restrict analysis to high confidence episodic recognition (rather than familiarity), and 2) ensure the inclusion of sufficient memory trials for robust estimation (e.g., Wimmer et al., 2014). Of the 174 participants who provided learning data, 165 also provided memory data (2 subjects discontinued the session, 7 had data loss from technical error). First, all trials where the participant responded that they were “guessing” were removed from analysis, but all “miss” trials were retained to be conservative. Then, participants were excluded if they had less than 120 signal trials (at least 2/3 of trials) after censoring (*N_excluded_* = 3). This resulted in memory data from 162 participants.

For all models, we compared linear and quadratic age predictors to examine whether age-related differences indicated an adolescent-specific (i.e., quadratic) or nonspecific (i.e., linear) effect (Somerville et al., 2013). Second-degree orthogonal polynomials of age were created using the poly function. Although generalized additive models are another data-driven, flexible way to examine the shape of age-related differences, we used polynomial models because the current implementation of generalized additive models cannot handle the complexity of the effects studied here (e.g., three-and four-way interactions). We performed F tests or likelihood ratio tests on the null, linear and quadratic age models to determine the best fit.

Mixed-effects logistic regression and beta-binomial models were fit using the glmmTMB package (Brooks et al., 2017). We performed regression model diagnostics with the performance and DHARMa packages (Hartig, 2024; Lüdecke et al., 2021). Standardized coefficients were computed using the effectsize package, conditional and marginal *R^2^* values were calculated using the performance package, and plots were created using sjPlot, ggeffects, and ggplot2 (Ben-Shachar et al., 2020; Lüdecke, 2018, 2023; Wickham, 2016).

### Analytic Approach

#### Learning Behavioral Analyses

Behavioral analyses were conducted to examine age-related differences in the effects of Pavlovian bias on instrumental learning. First, we submitted trial-by-trial learning accuracy to a set of nested mixed-effects logistic regression models with random intercepts for participants. We performed likelihood ratio tests to determine the best-fitting model with the stepwise addition of the following hypothesis-driven fixed effects: mean-centered trial number, action (go, ref: no-go), valence (win, ref: avoid loss), linear age, and quadratic age, as well as all possible interactions (Table S1). Then, we performed likelihood ratio tests to determine whether iteratively removing non-significant, highest-order interactions resulted in significantly decreased model fit. We first removed the non-significant, four-way interaction of Trial*Action* Valence*Age then the non-significant, three-way interaction of Trial*Valence*Age. This parsimonious model had no significant decrease in model fit compared to the models with these higher-order interactions (Table S2). Results from the parsimonious model are reported in Table 1, and results from the complex model including the higher-order interaction terms are presented in Table S3. Wald tests were used to compute *p* values and determine the statistical significance of the coefficients.

Next, we conducted post-hoc analyses based on prior developmental studies of this paradigm (Betts et al., 2020; Moutoussis et al., 2018; Raab & Hartley, 2020). We used paired t-tests comparing the Pavlovian-congruent and incongruent condition within each action condition (i.e., Go to Win vs. Go to Avoid Loss) to characterize Pavlovian influence overall (i.e., not associated with age). We also used paired t-tests of accuracy in the first and last task blocks within each trial type to determine the extent to which participants learned the optimal responses over time.

We investigated age-related differences in overall performance and performance in each action-valence condition by submitting accuracy to beta-binomial regressions of linear and quadratic age. Beta-binomial models were selected to account for overdispersion (Ferrari & Comelli, 2016), and simulated posterior predictive checks indicated that beta-binomial models captured the observed data better than linear and binomial regression models (Fig. S6). We also examined age-related differences in reaction time to Pavlovian-congruent “Go to Win” and Pavlovian-incongruent “Go to Avoid Loss” using a mixed-effects gamma regression, which accounts for the zero-bounded, right-skewed shape of RT. Making a response is the correct action for both trial types, so if participants are relatively faster during the “Go to Win” trials, this would suggest that the potential to gain reward or Pavlovian-congruency facilitates faster responding. For this model, we included random intercepts for participants and performed likelihood ratio tests to determine the best-fitting model with the stepwise addition of the following fixed effects: valence (win, ref: avoid loss), linear age, and quadratic age, as well as all possible interactions.

Motivated by a previous study that found attenuated effects of Pavlovian bias in adolescence (Raab & Hartley, 2020), we also calculated a behavioral metric of Pavlovian bias and submitted it to regressions of linear and quadratic age:

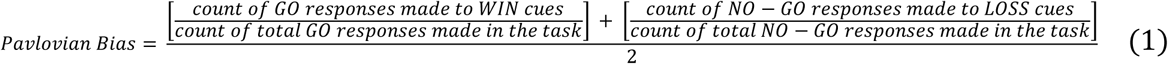

#### RL Models

Next, we fit a set of RL models to participants’ choice data. These computational models provide a formalized approach to test the latent psychological processes contributing to learning. For example, we can test whether age-related differences in choice behavior are attributable to differences in how quickly an individual learns from reinforcement (i.e., differences in the learning rate), or in how sensitive an individual is to the outcome (i.e., differences in reinforcement sensitivity). These models track the values of each potential action (go, no-go) for each stimulus across reinforcement. The basic model included a Rescorla-Wagner update policy for the action value Q(a,s) on each trial t (Barto & Sutton, 1982; Rescorla & Solomon, 1967; Sutton & Barto, 1981):

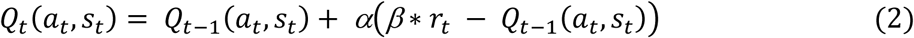

where a_t_, s_t_, and r_t_ represent the action taken (go: *1* or no-go: *0*), the state (e.g., Go to Win: *1, 2, 3, 4*), and the outcome (either *-.25, 0, .25*) for the trial, respectively; α is the learning rate (bounded from 0 to 1), and β is the reinforcement sensitivity (bounded from 0 to +∞). The action values were transformed into probabilities of selecting an action given the current state through a squashed softmax function:

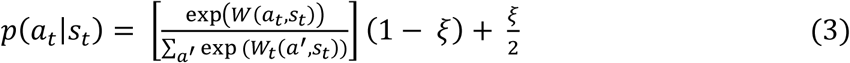

where ξ is the lapse rate parameter, which was free to vary between 0 and 1.

We compared more complex models with two learning rates (α_v_) or two reinforcement sensitivities for the Valence conditions (β_v_) to test whether participants learned differently from or were differently sensitive to gains and losses. For some models, we augmented the value of “go” with a bias *b* towards pressing the button (“go bias”):

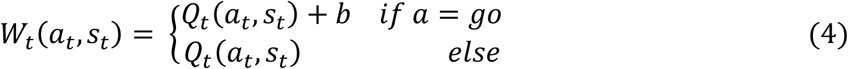

Additionally, we included a Pavlovian bias parameter *π* (bounded from 0 to +∞) that was multiplied by the Pavlovian stimulus value V_t_, which is updated every trial through reinforcement disregarding the choice:

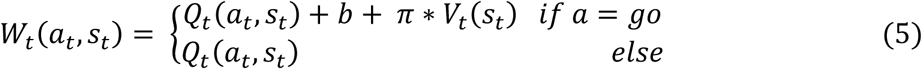

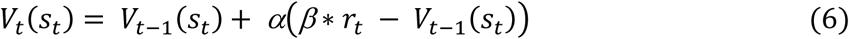

The Pavlovian stimulus value V_t_ is positive for Win conditions, as participants receive win and neutral outcomes in these conditions, and is negative for Avoid Loss conditions, as they receive neutral and loss outcomes in these conditions. Thus, the Pavlovian bias parameter promotes the tendency to go in proportion to the positive value of Win stimuli and to no-go in proportion to the negative value of Loss stimuli. We also tested models that included two Pavlovian bias parameters: we tested a Valence condition model (*π*_*v*_) that accounted for whether the state was to Win or to Avoid Loss, and a Pavlovian-congruency model (*π*_*c*_) that accounted for whether the state was congruent or incongruent.

Model fitting and parameter estimation were performed using the emfit package developed by Quentin Huys, 2018 for Matlab, and was downloaded from Huys’ GitHub (public domain). This package conducts a hierarchical model-fitting procedure that uses maximum likelihood to estimate group-level and individual-level parameter distributions (Guitart-Masip et al., 2012; Huys et al., 2011). On each iteration, the model is fit to the participant’s choice data to estimate individual-level parameters, which then contribute to group-level estimates for each parameter, which are then used as priors for the individual-level estimates for the next iteration. All participants were included in one group for estimation, which is a conservative approach to testing age-related differences as individual estimates are constrained towards the group mean. We determined the relatively best-fitting model using the integrated Bayesian Information Criterion (iBIC), as is standard for this model-fitting procedure (Table S4; Betts et al., 2020; Guitart-Masip et al., 2012; Huys et al., 2011). The iBIC balances fit and parsimony by penalizing for the number of model parameters. To ensure the winning model captured participant choice behavior well, we simulated new choice data using the parameter estimates from each individual and verified that it qualitatively reproduced participant choice behavior (see Figs. S8-S9 for model validation and parameter recovery). We constructed regressions to test whether each parameter was linearly or quadratically related to age, which could inform whether the adolescent peak in performance was driven by a specific cognitive computation, such as gain sensitivity.

#### Memory Behavioral Analyses

We examined age-related differences in overall memory accuracy by comparing models of *d’* with linear and quadratic age as predictors. We calculated the participants’ overall sensitivity (*d’*) using signal detection techniques in the psycho package (*d’ = z(hits / (hits + misses)) – z(false alarms / (false alarms + correct rejects))*; Makowski, 2018; Stanislaw & Todorov, 1999). This measure of sensitivity corrects the hit rate for participants’ tendency to identify an image as “Old.” We also tested whether participants of different ages showed a reward memory bias, by calculating *d’* separately for images shown when the participants could gain a reward and did gain, and when they could gain a reward but did not gain (i.e., for images appearing in rewarded and not rewarded Win trials). We submitted *d’* to nested mixed-effects models and used likelihood ratio tests to determine the best-fitting model with the stepwise addition of the following fixed effects: reward (gain, ref: no gain), linear age, and quadratic age, and reward*age interactions, with random intercepts for participants.

Then, we used mixed-effects logistic regression models to address our key research question about age-related differences in the strength of the association between prediction error and memory. First, we computed trial-by-trial prediction errors by applying the learning model and group-level parameters to the participants’ sequences of stimuli, choices, and outcomes. Following previous studies, we used group-level parameter estimates, because they result in less noise in the prediction errors than individual-level estimates (Eldar et al., 2016; Hauser et al., 2017; Seymour et al., 2012). Second-degree orthogonal polynomials of prediction error were created using the poly function to partial out the effects of signed and unsigned prediction errors (i.e., linear and quadratic effects, respectively). We performed likelihood ratio tests to determine the best-fitting model for trial-by-trial memory accuracy with the stepwise addition of the following fixed effects and their interactions: linear and quadratic prediction error, and linear and quadratic age (Table S5). All models included random intercepts for participants. To understand the model results, we plotted predicted results at representative ages (i.e., 10, 15, 20, and 25-years-old) using the ggeffects package (Lüdecke, 2018).

## Acknowledgements

The authors wish to thank Michael J. Frank, Nina Rouhani, and Patrick Mair for helpful discussions; Quentin Huys for making his code publicly available; and the Research Assistants in the Learning and Brain Development Lab at Northeastern for assistance with participant recruitment and testing. We are grateful to the participants and families who took part in this study. This research was supported in part by the National Science Foundation (DGE-1938052 to H.M.H. and Career Award 2443141 to J.Y.D.).

## SUPPLEMENTAL INFORMATION

**Figure S1.**
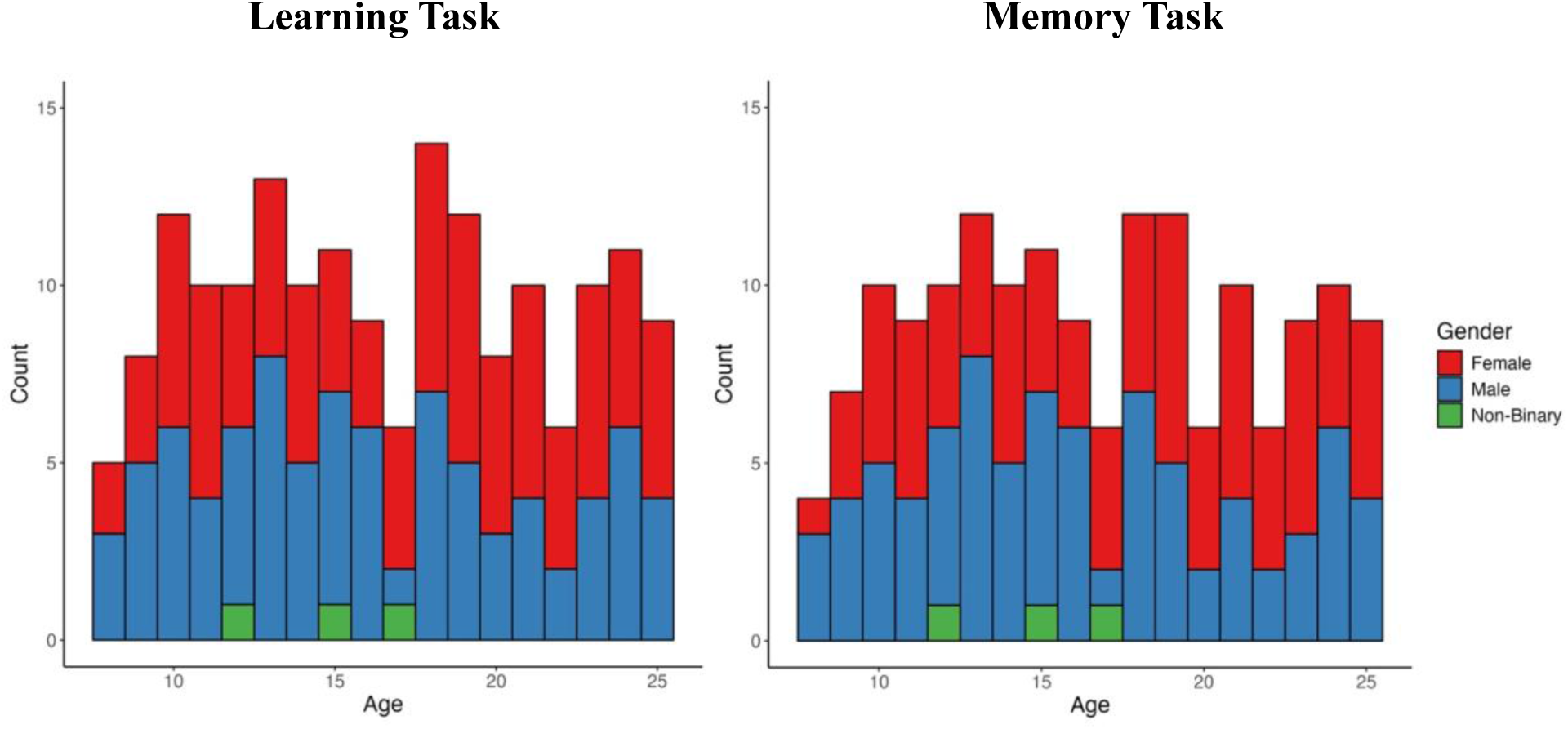
Sample Demographics. The left plot shows the sample demographics for people who completed the learning task (*N* = 174). A subset of participants also completed the memory task with adequate performance, shown on the right (*N* = 162). Both samples have good representation across the age range.

### Learning Behavioral Analyses

**Table S1.** Model Comparison for Nested Learning Models. Models increase in complexity starting with a null model. Predictors are added in order of largest expected effect. Likelihood ratio tests indicate that the best-fitting model includes all predictors of interest, the FE Quadratic Age model. RE = random effect. FE = fixed effect.

| Sampling Units |  | N total observations: 31320<br>N Subjects: 174; N Trials: 180 |  |  |  |  |  |  |
| --- | --- | --- | --- | --- | --- | --- | --- | --- |
| Model Specification | Nested Model | Fixed Effects Added | Random Effects | AIC | BIC | LL | LRT df | LRT $X^2$ |
| RE Only | - | - | Subject intercepts | 40617 | 40633 | -20306 | - | - |
| FE Trial | RE Only | Trial | “” | 40416 | 40441 | -20205 | 1 | 202.31* |
| FE Action | FE Trial | Trial x Action | “” | 38530 | 38572 | -19260 | 2 | 1890.42* |
| FE Valence | FE Action | Trial x Action x Reward | “” | 37715 | 37791 | -18849 | 4 | 822.61* |
| FE Linear Age | FE Valence | Trial x Action x Reward x Age | “” | 37676 | 37818 | -18821 | 8 | 55.83* |
| FE Quadratic Age | FE Linear Age | Trial x Action x Reward x Age + Age<sup>2</sup> | “” | 37580 | 37788 | -18765 | 8 | 111.95* |

**Table S2.** Model Comparison for Reduced Learning Models. Highest-order non-significant interactions are removed from the model before testing for degraded model fit. Likelihood ratio tests indicate that including the non-significant, higher-order interactions do not result in improved model fit, so we report results from the FE Quadratic Age – Parsimonious Model.

| Model Specification | Nested Model | Fixed Effects Added | Random Effects | AIC | BIC | LL | LRT df | LRT $X^2$ |
| --- | --- | --- | --- | --- | --- | --- | --- | --- |
| **FE Quadratic Age - Parsimonious** |  | - | **Subject intercepts** | **37577** | **37752** | **-18768** | - | - |
| FE Quadratic Age - Reduced | FE Quadratic Age - Parsimonious | Trial x Reward x Age + Age<sup>2</sup> | “” | 37578 | 37771 | -18766 | 2 | 2.60 |
| FE Quadratic Age | FE Quadratic Age - Reduced | Trial x Action x Reward x Age + Age<sup>2</sup> | “” | 37580 | 37788 | -18765 | 2 | 2.84 |

**Figure S2.**
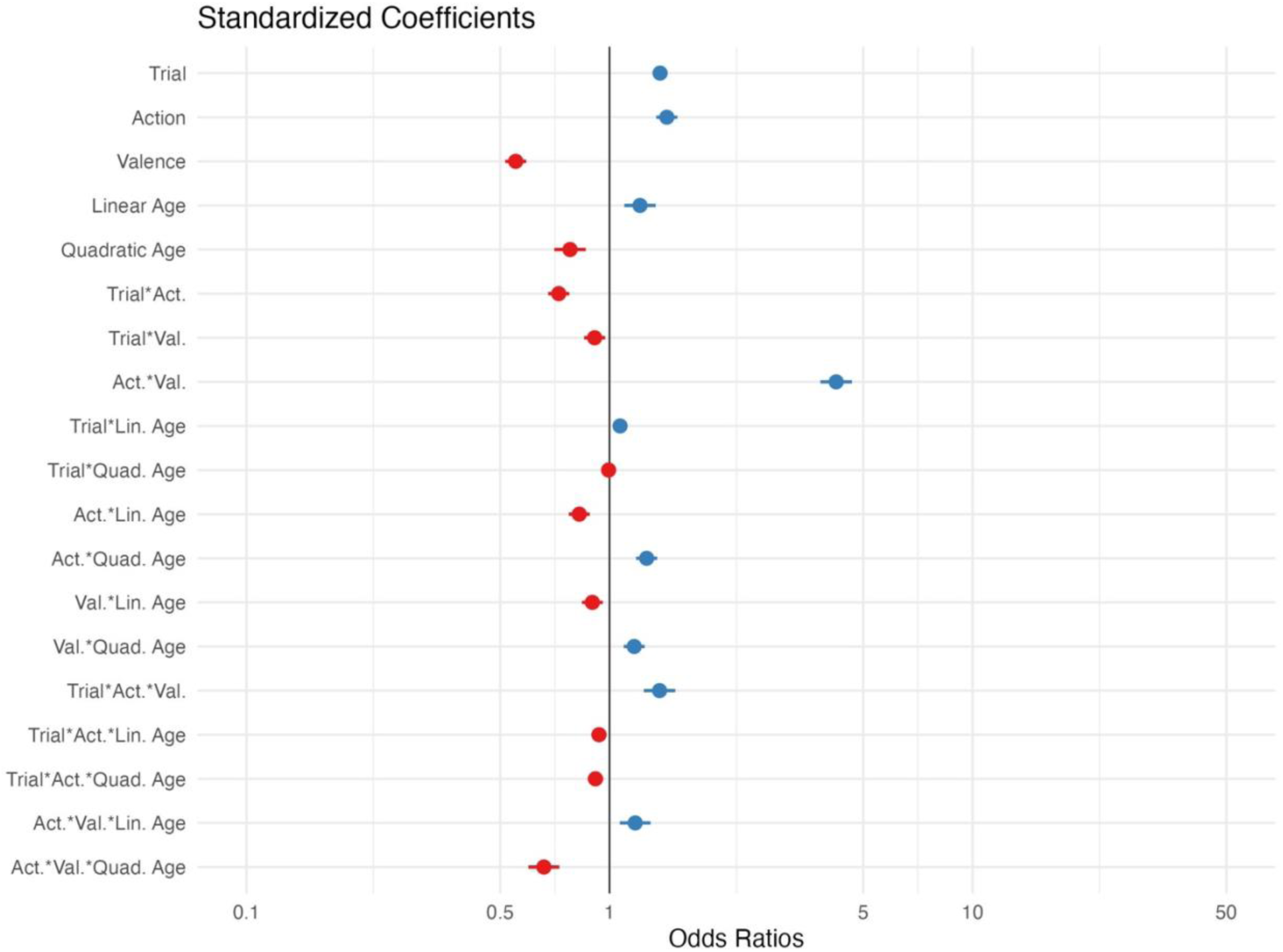
Best-Fitting Model Standardized Odds Ratios. Odds ratios less than one correspond to “negative” coefficients (i.e., the odds of a correct response decrease with an additional unit of the predictor), and ratios greater than one correspond to “positive” coefficients (i.e., the odds of a correct response increase with an additional unit of the predictor).

**Table S3.** Results from the More Complex Model. The results of the more complex model (FE Quadratic Age) include the non-significant, higher-order interactions that are removed from the Parsimonious model. The odds ratios are similar to those presented in the main text (Table 1) estimated by the parsimonious model.

| Fixed Effects |  |  |  |  |
| --- | --- | --- | --- | --- |
|  | Odds Ratio | 95% CI | Z-value | p-value |
| Intercept | 1.31 | 1.18-1.45 |  |  |
| **Trial** | **1.38** | **1.31-1.44** | **13.20** | **< .001** |
| **Action: Go (v. No Go)** | **1.44** | **1.34-1.54** | **10.62** | **< .001** |
| **Valence: Win (v. Avoid Loss)** | **0.55** | **0.52-0.59** | **-17.48** | **< .001** |
| **Age** | **1.21** | **1.10-1.34** | **3.80** | **< .001** |
| **Age<sup>2</sup>** | **0.78** | **0.70-0.86** | **-4.92** | **< .001** |
| **Trial x Action** | **0.72** | **0.68-0.77** | **-9.41** | **< .001** |
| **Trial x Valence** | **0.91** | **0.85-0.97** | **-2.75** | **.006** |
| **Action x Valence** | **4.23** | **3.83-4.68** | **28.13** | **< .001** |
| **Trial x Age** | **1.06** | **1.01-1.11** | **2.22** | **.026** |
| Trial x Age<sup>2</sup> | 0.99 | 0.94-1.04 | -0.39 | .700 |
| **Action x Age** | **0.83** | **0.77-0.88** | **-5.64** | **< .001** |
| **Action x Age<sup>2</sup>** | **1.27** | **1.18-1.35** | **6.89** | **< .001** |
| **Valence x Age** | **0.90** | **0.84-0.96** | **-3.23** | **.001** |
| **Valence x Age<sup>2</sup>** | **1.17** | **1.09-1.25** | **4.62** | **< .001** |
| **Trial x Action x Valence** | **1.38** | **1.25-1.53** | **6.34** | **< .001** |
| Trial x Action x Age | 0.94 | 0.88-1.00 | -1.86 | .064 |
| Trial x Action x Age<sup>2</sup> | 0.95 | 0.89-1.01 | -1.53 | .125 |
| Trial x Valence x Age | 1.03 | 0.96-1.10 | 0.79 | .431 |
| Trial x Valence x Age<sup>2</sup> | 1.01 | 0.94-1.08 | 0.20 | .838 |
| **Action x Valence x Age** | **1.18** | **1.07-1.30** | **3.29** | **.001** |
| **Action x Valence x Age<sup>2</sup>** | **0.66** | **0.60-0.73** | **-8.35** | **< .001** |
| Trial x Action x Valence x Age | 0.99 | 0.90-1.10 | -0.13 | .895 |
| Trial x Action x Valence x Age<sup>2</sup> | 0.92 | 0.83-1.01 | -1.68 | .093 |
| Random Effects |  |  |  |  |
|  | Variance | Standard Deviation |  |  |
| Subject (Intercept) | 0.35 | 0.59 |  |  |
| Model Equation: Accuracy ~ Trial*Action*Valence*poly(Age,2) + (1\|Subject) |  |  |  |  |

**Figure S3.**
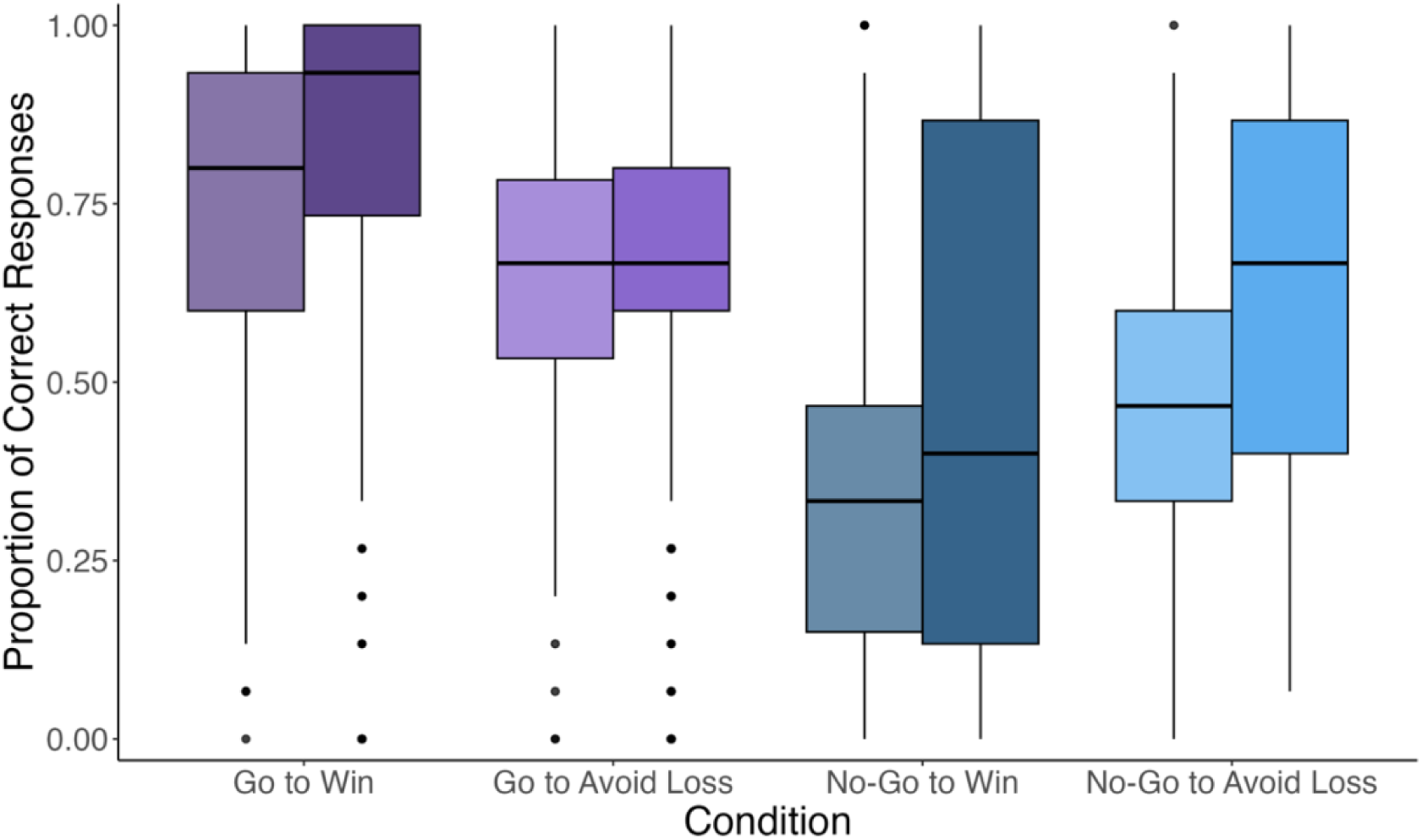
Learning within each Condition. The influence of Pavlovian bias on instrumental learning was evident at the group level, whereby participants had higher accuracy in the Pavlovian-congruent than incongruent conditions. This is generally consistent with prior literature employing this task, whether in adults only, across development, or throughout the lifespan. Participants performed better in the Go to Win than the Go to Avoid Loss condition (*M_Difference_* = .15, 95% CI = [.12, .18], *t*(173) = 10.40, *p* < .001) and in the No-Go to Avoid Loss than the No-Go to Win condition (*M_Difference_* = .13, 95% CI = [.09, .17], *t*(173) = 6.47, *p* < .001). Further, paired t-tests of accuracy in the first and last task blocks indicated that accuracy increased across the task in the Go to Win (*M_Difference_* = .08, 95% CI = [.04, .12], *t*(173) = 3.68, *p* < .001), No-Go to Win (*M_Difference_* = .11, 95% CI = [.06, .16], *t*(173) = 4.13, *p* < .001), and No-Go To Avoid Loss (*M_Difference_* = .16, 95% CI = [.12, .20], *t*(173) = 7.39, *p* < .001) conditions, but not in the Go to Avoid Loss (*M_Difference_* = .01, 95% CI = [−.03, .04], *t*(173) = 0.29, *p* = .77) condition. Light and dark bars represent accuracy in the first and last blocks of the task (averaged over 15 trials each), respectively, reflecting learning over time. The probabilistic frequency of the optimal outcome was set to 80%.

**Figure S4.**
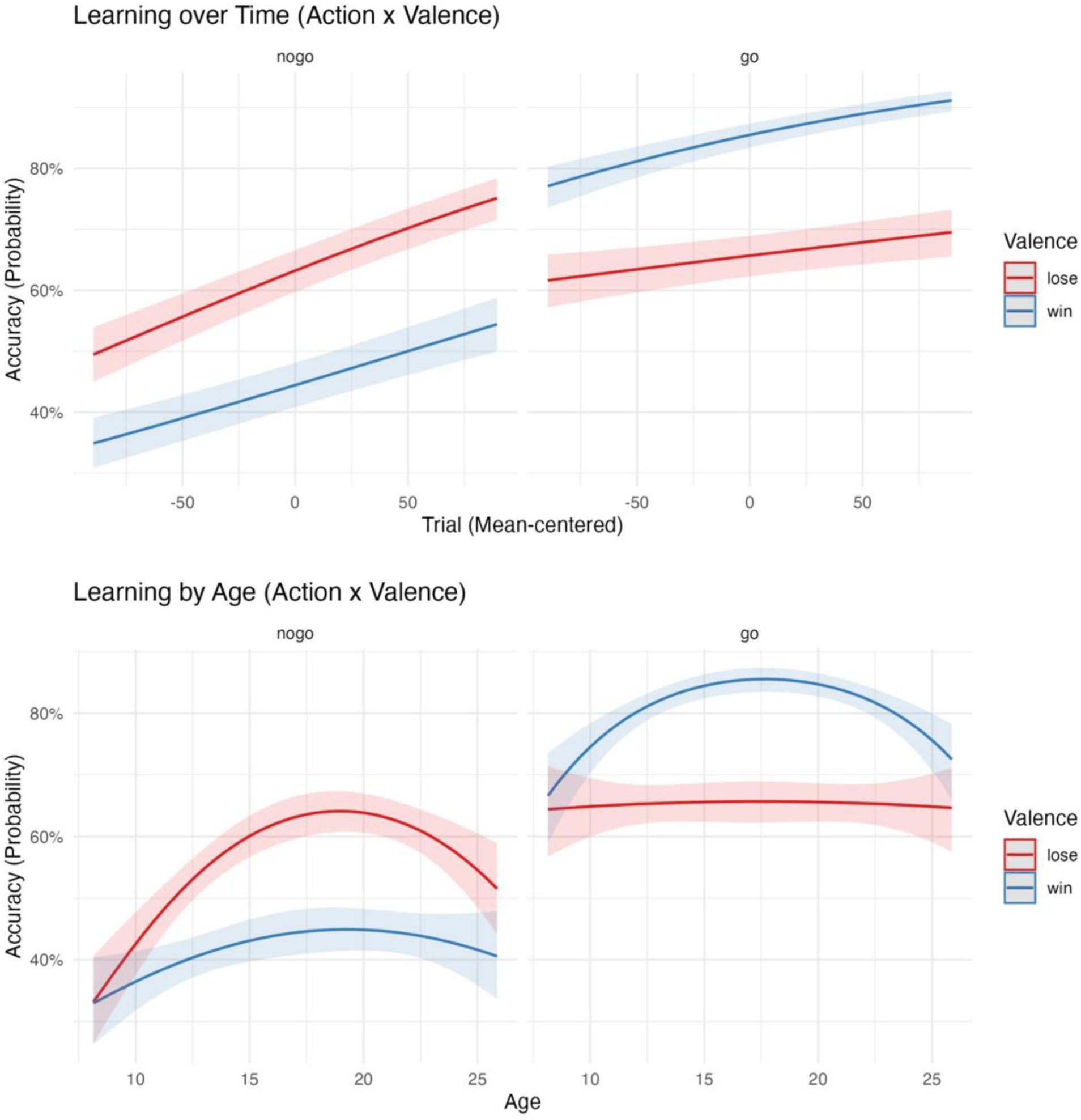
Effects Plots of Key Interactions. The first plot shows the model predictions for the Action*Valence*Trial Number interaction (at the sample mean age 17.09). The slopes of the lines indicate the rate of learning across the task for each trial type. The second plot shows the model predictions for the Action*Valence*Age interaction (at the mean trial). The adolescent peak in performance is more prominent in the Pavlovian-congruent conditions (Go to Win and No Go to Avoid Loss) than the Pavlovian-incongruent conditions. Shaded regions show the 95% confidence intervals.

**Figure S5.**
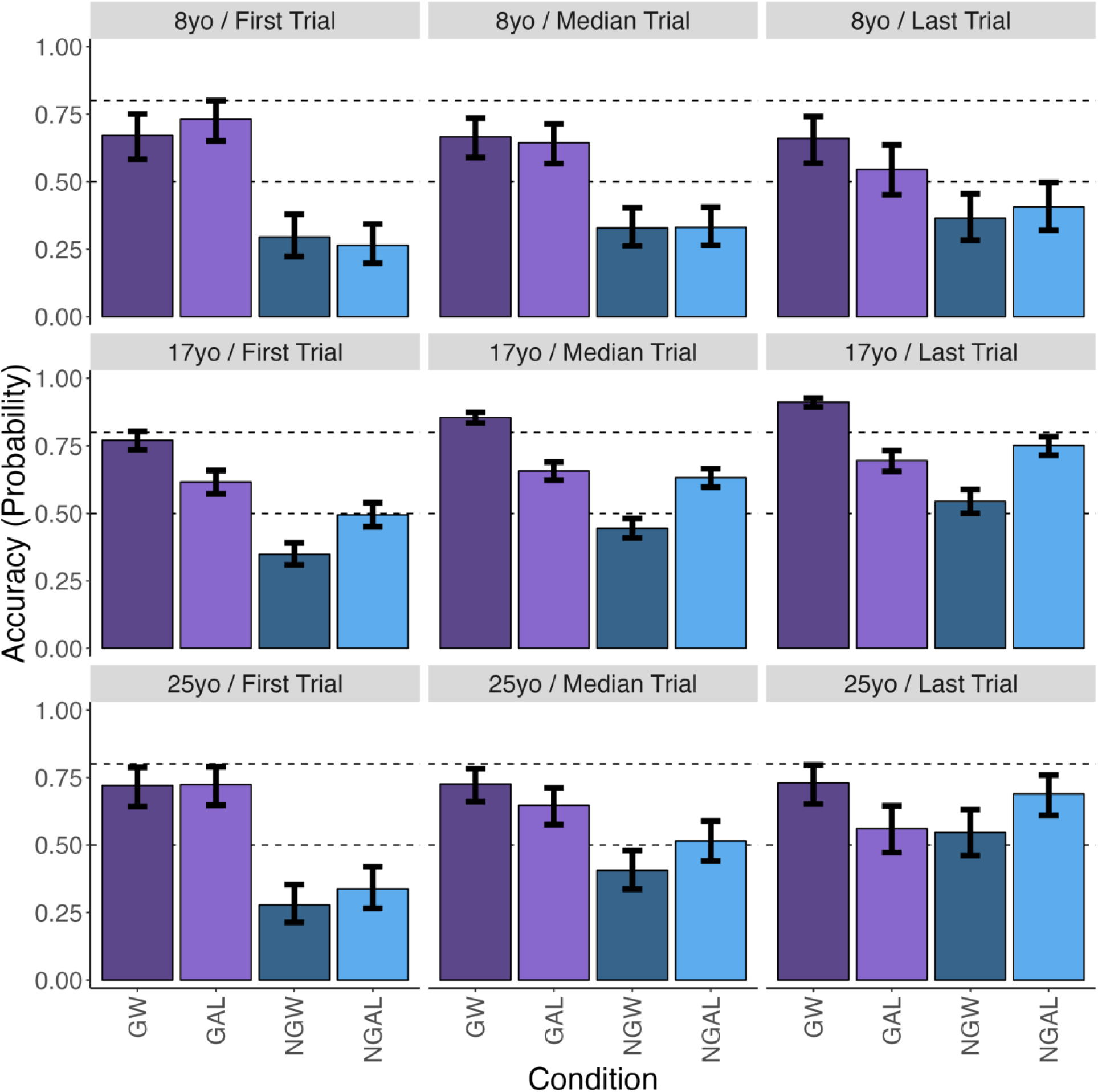
Predicted Learning Accuracy at Different Ages and Trials. The bars show the predicted probability of a correct response at the first, median, and last trials for participants aged 8.15 (sample minimum), 17.09 (sample mean), and 25.86 (sample maximum) years old. The “17yo / Last Trial” plot (middle row, right column) is the same as Figure 2a in the main text. The error bars represent 95% confidence intervals. Pavlovian-congruent conditions are outer bars (GW & NGAL) and Pavlovian-incongruent are inner bars (GAL & NGW). Dashed lines are at 50% (chance responding to either go or no-go) and 80% (reflecting the probabilistic frequency of best outcome per condition in this task).

**Figure S6.**
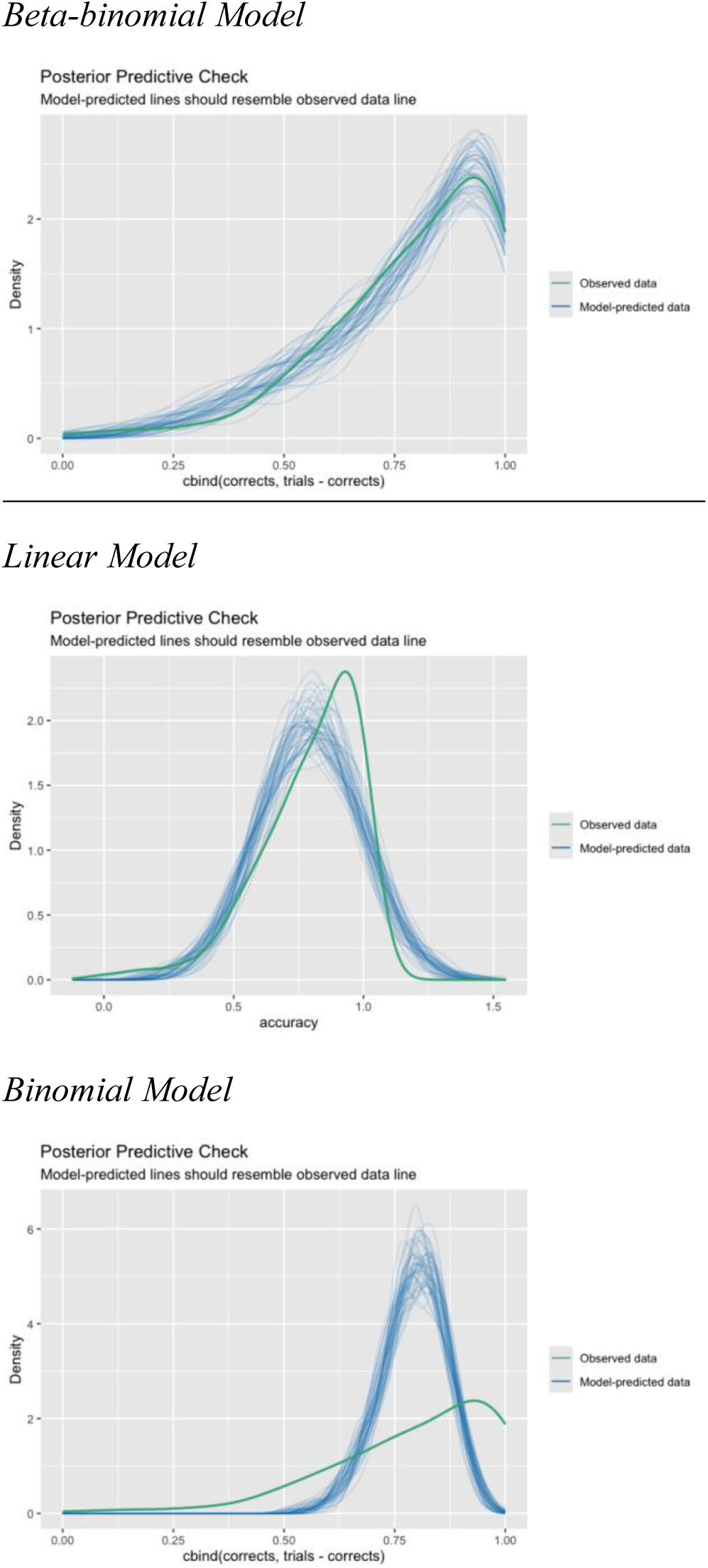
Comparing Linear, Binomial, and Beta-binomial Regression Models for “Go to Win.” Simulated data from the beta-binomial model (top) were more similar to the observed data (i.e., the blue lines overlap the green more closely) than simulated data from linear (middle) and binomial (bottom) models. These posterior predictive checks were conducted using the performance package (Lüdecke et al., 2021).

**Figure S7.**
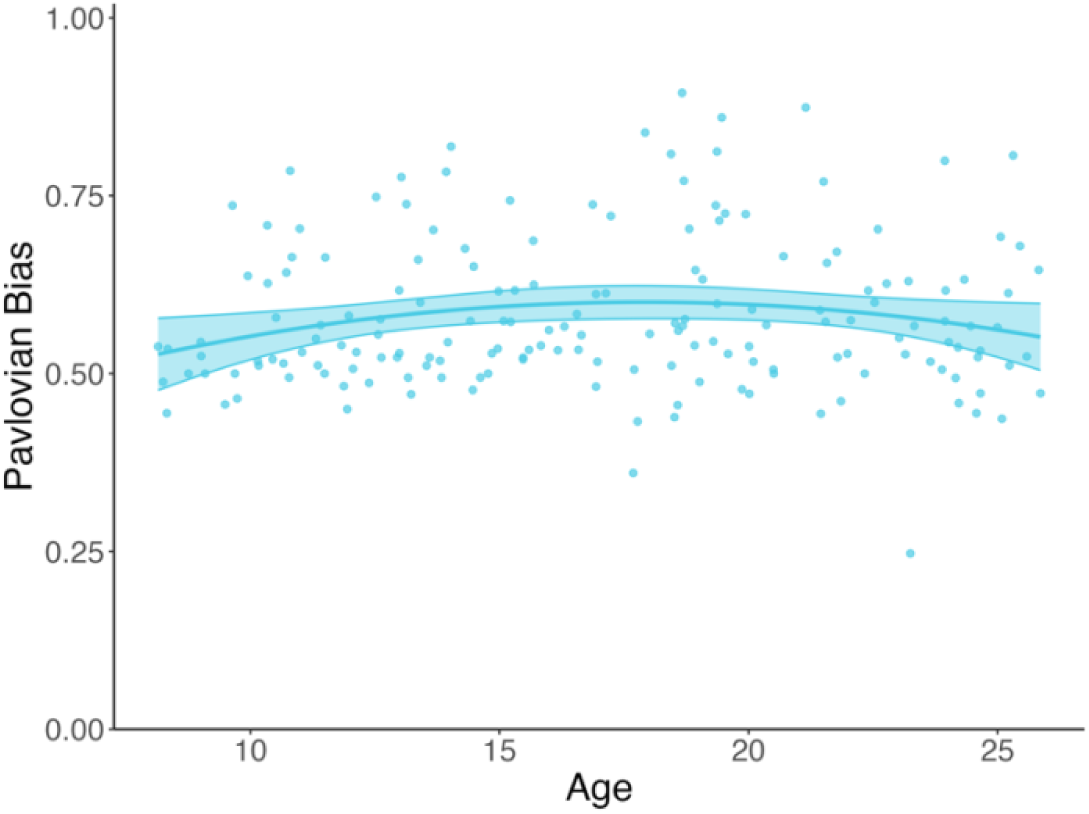
Age-related Differences in Behavioral Pavlovian Bias Scores. The behavioral metric of Pavlovian bias had a shallow quadratic association with age, with an adolescent boost in the score (Quadratic vs. Linear Model: *F*(1,171) = 4.90, *p* < .05). The line represents model predictions across age, shaded regions show 95% CIs, and points show raw data.

### Reinforcement Learning (RL) Modeling Information

**Table S4.**
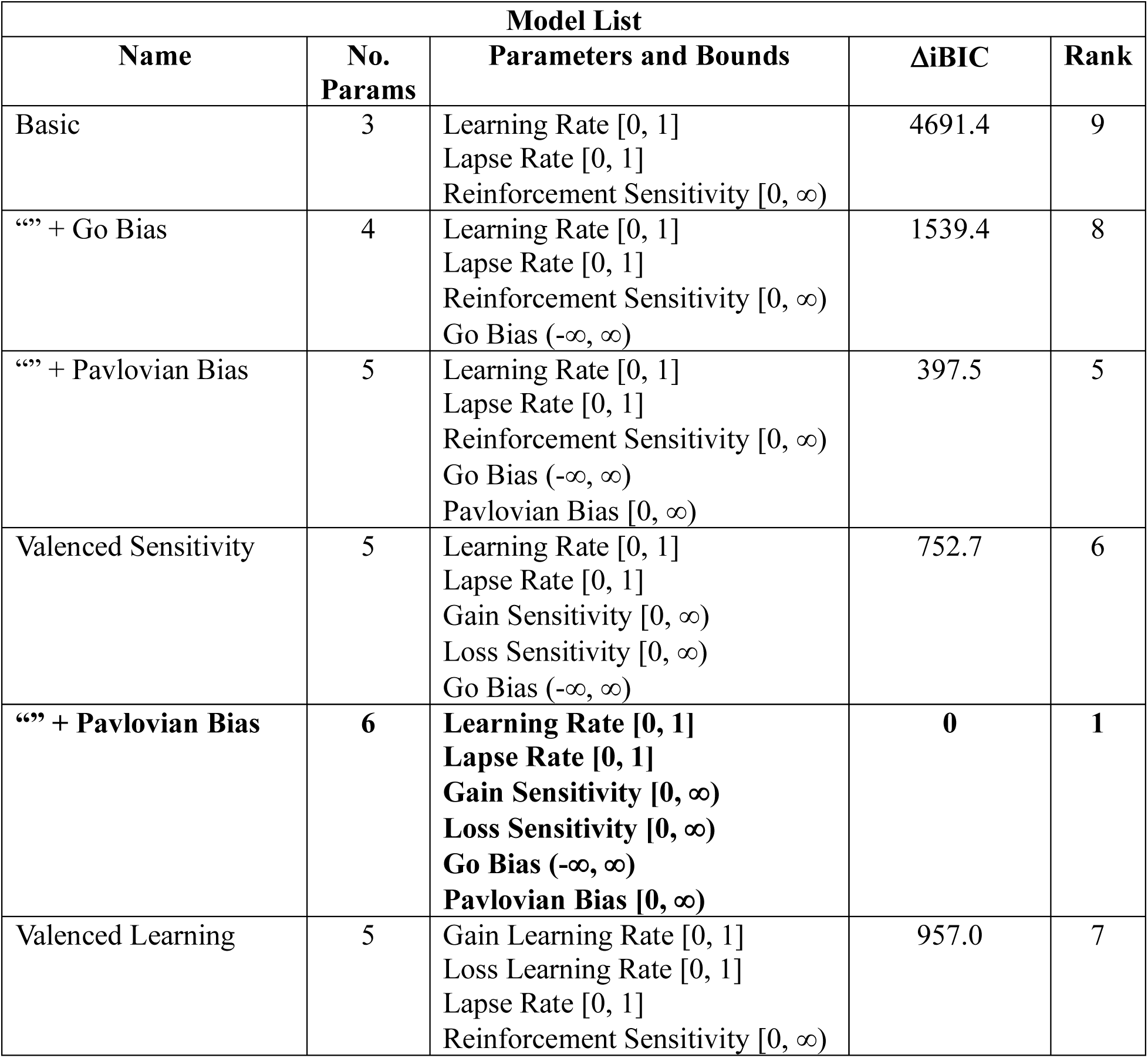
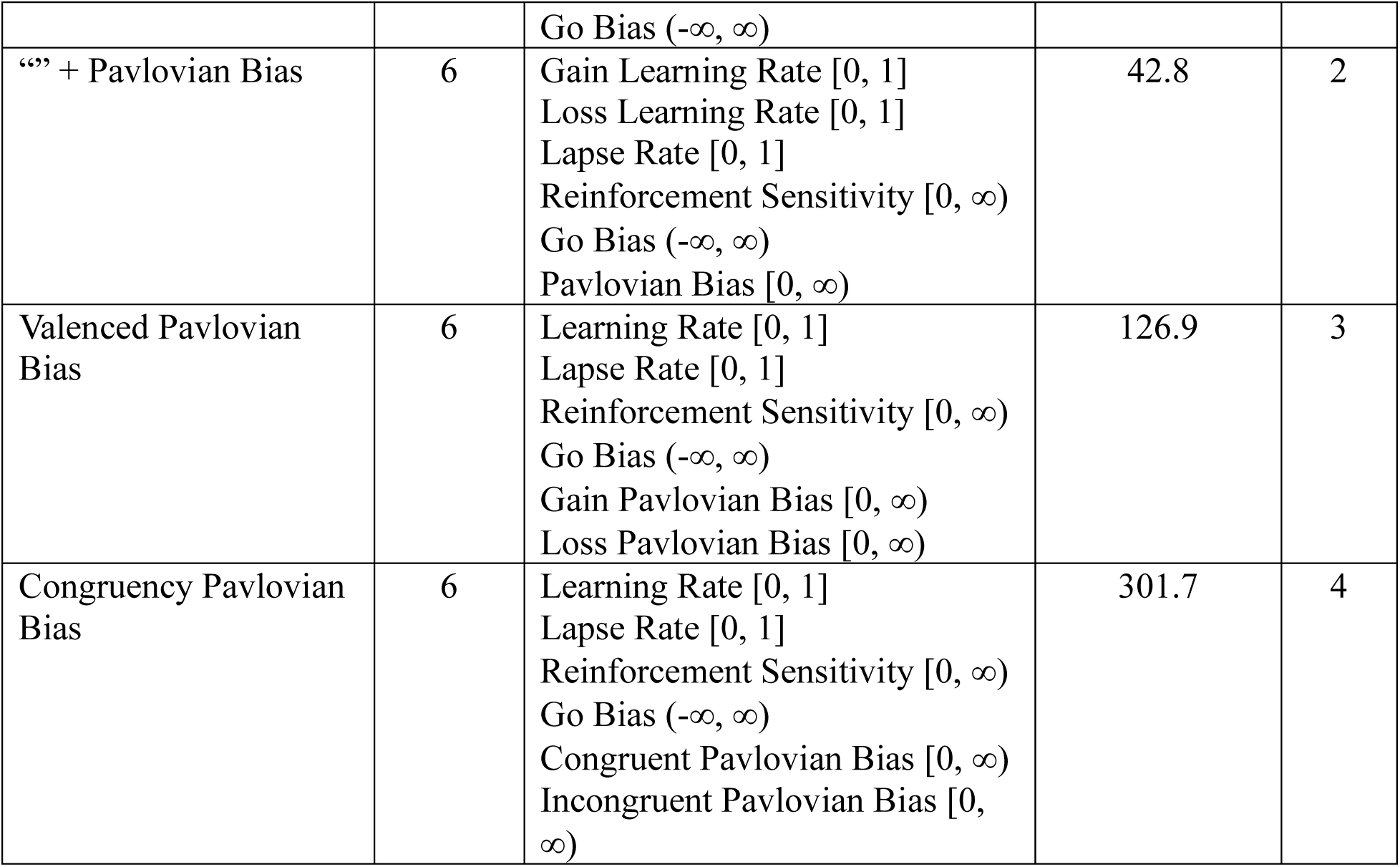
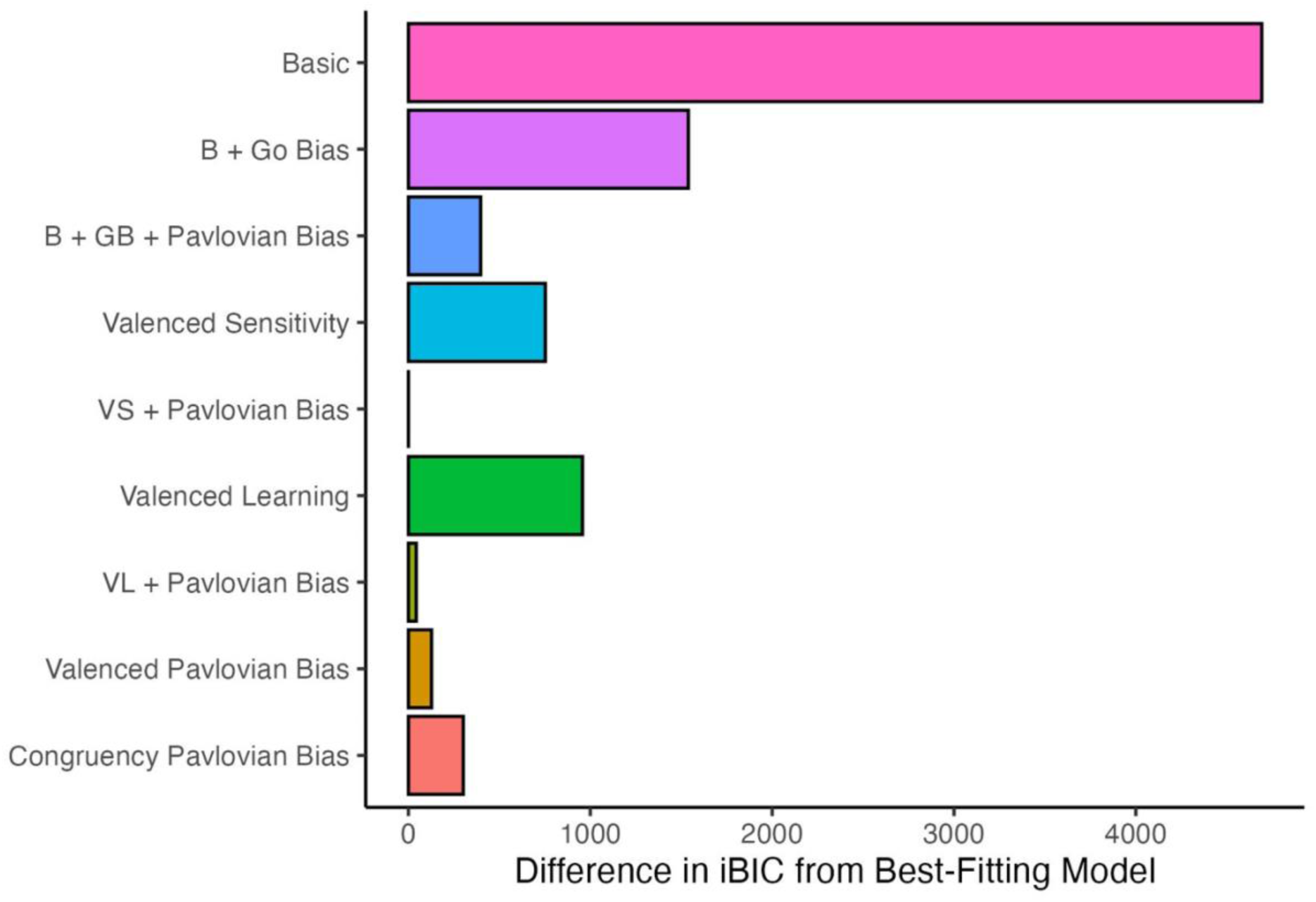
Model List. We tested a set of nine RL models, which are described in this table. The winning model (5^th^ row, bolded) included a learning rate, lapse rate, gain and loss sensitivities, go bias, and Pavlovian bias parameters. To evaluate which of the alternative reinforcement learning models had the most explanatory power, we computed an integrated Bayes Information Criterion (iBIC) for the log likelihood estimated from the best-fit iteration for each model. The Bayes Information Criterion (BIC, Schwarz, 1978) for each subject was summed for each model separately, with the lowest value indicating the best model. Following Cavanagh et al., 2013, differences in iBIC between alternative models greater than 20 are highly suggestive of a meaningful difference (Kass & Raftery, 1995). The difference here between the best-fit model and the model with the next lowest iBIC was 42.8. This model also had the same number of fit parameters (6).

**Figure S8.**
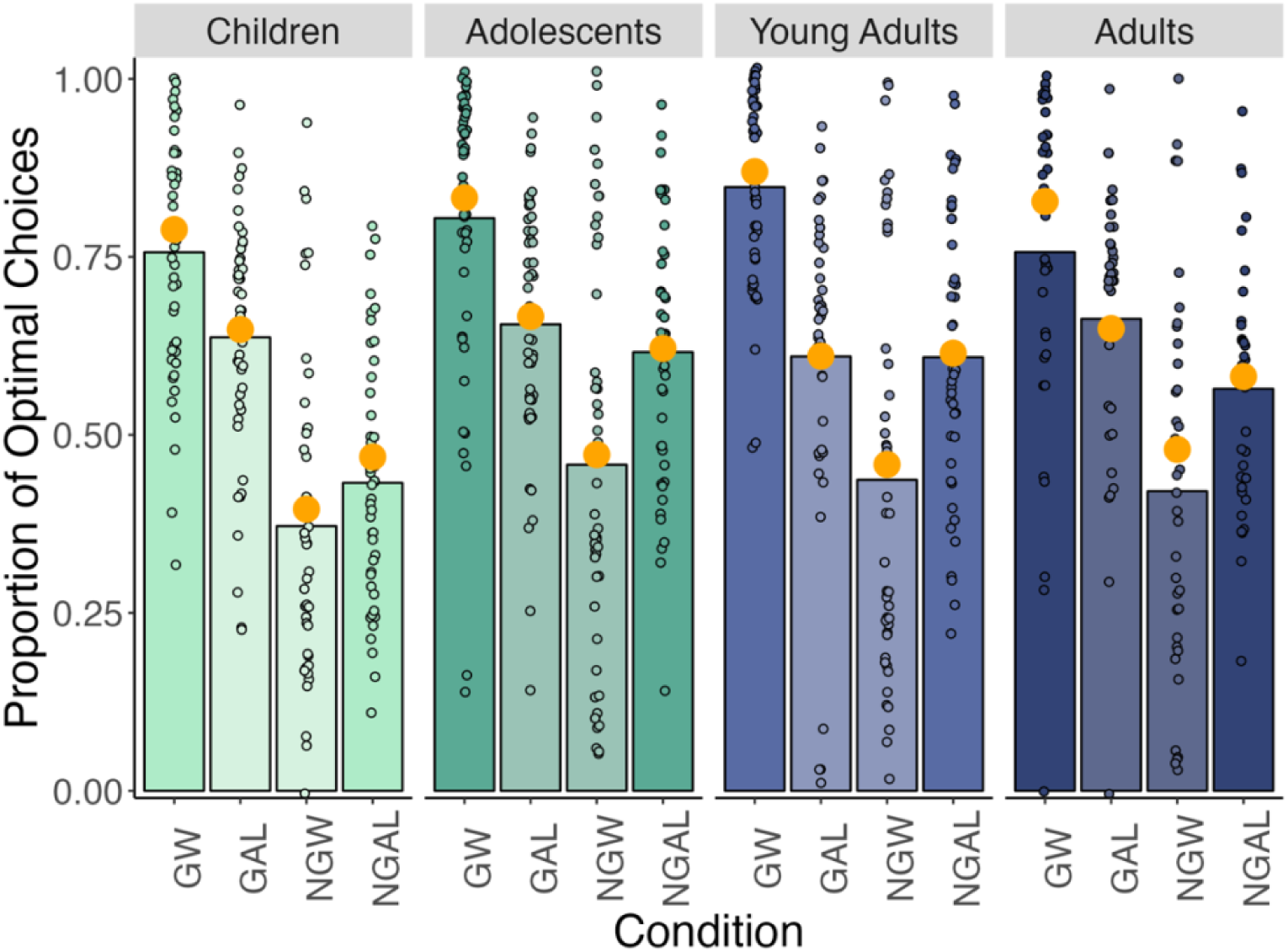
Model Validation. The bars represent mean accuracy within each age group (Children: 8 – 12.99; Adolescents: 13 – 17.99; Young Adults: 18 – 21.99; Adults: 22 – 25.99) for each trial type from the participants’ choice data (dark: Pavlovian-congruent, light: Pavlovian-incongruent). Points show each participants’ accuracy. The large orange points are the mean accuracy for each age group generated from the winning RL model, computed as the average from 100 simulations of the task using each set of individual-level parameter estimates (*N* = 174). The model recapitulates participants’ choice behavior well, although there is some deviation for the adults in the Win conditions.

**Figure S9.**
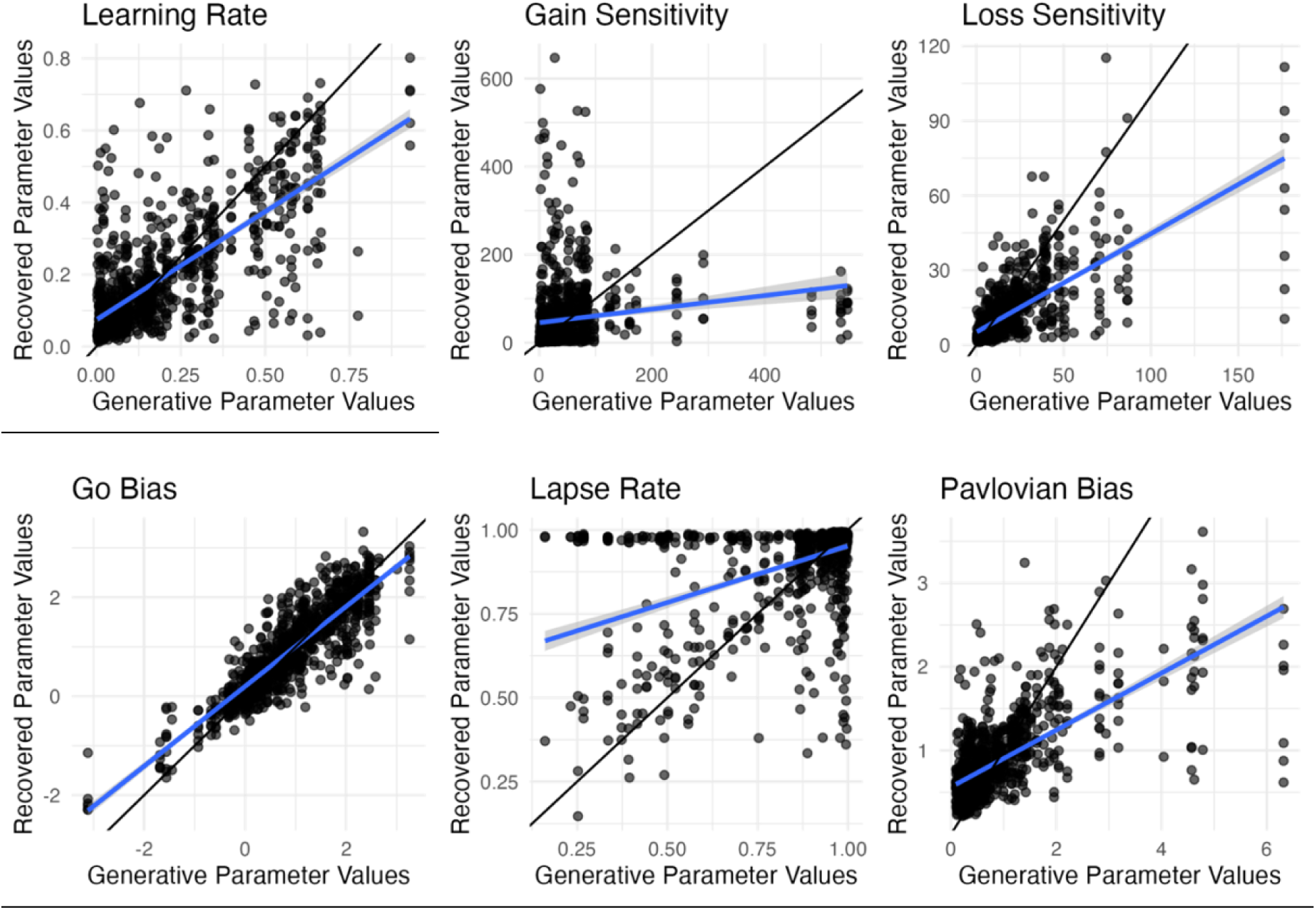
Parameter Recovery. Parameter recovery was assessed by generating data for 1000 subjects, using bootstrapped samples of the individual parameter solutions. The best-fitting model was fit to these data, and the estimates of the recovered parameters were correlated with the generative parameters. Parameter recovery was adequate (Spearman’s rhos = .42 - .87, with lapse rate and go bias as the best and worst recovered, respectively), but as the expectation maximization approach tends to constrain estimates towards the group mean, some of the more extreme values were poorly recovered and should be interpreted cautiously. Each point represents one of the 1000 estimates. The black line represents the “ideal” linear association between the estimates, and the blue line represents the estimated linear association.

**Figure S10.**
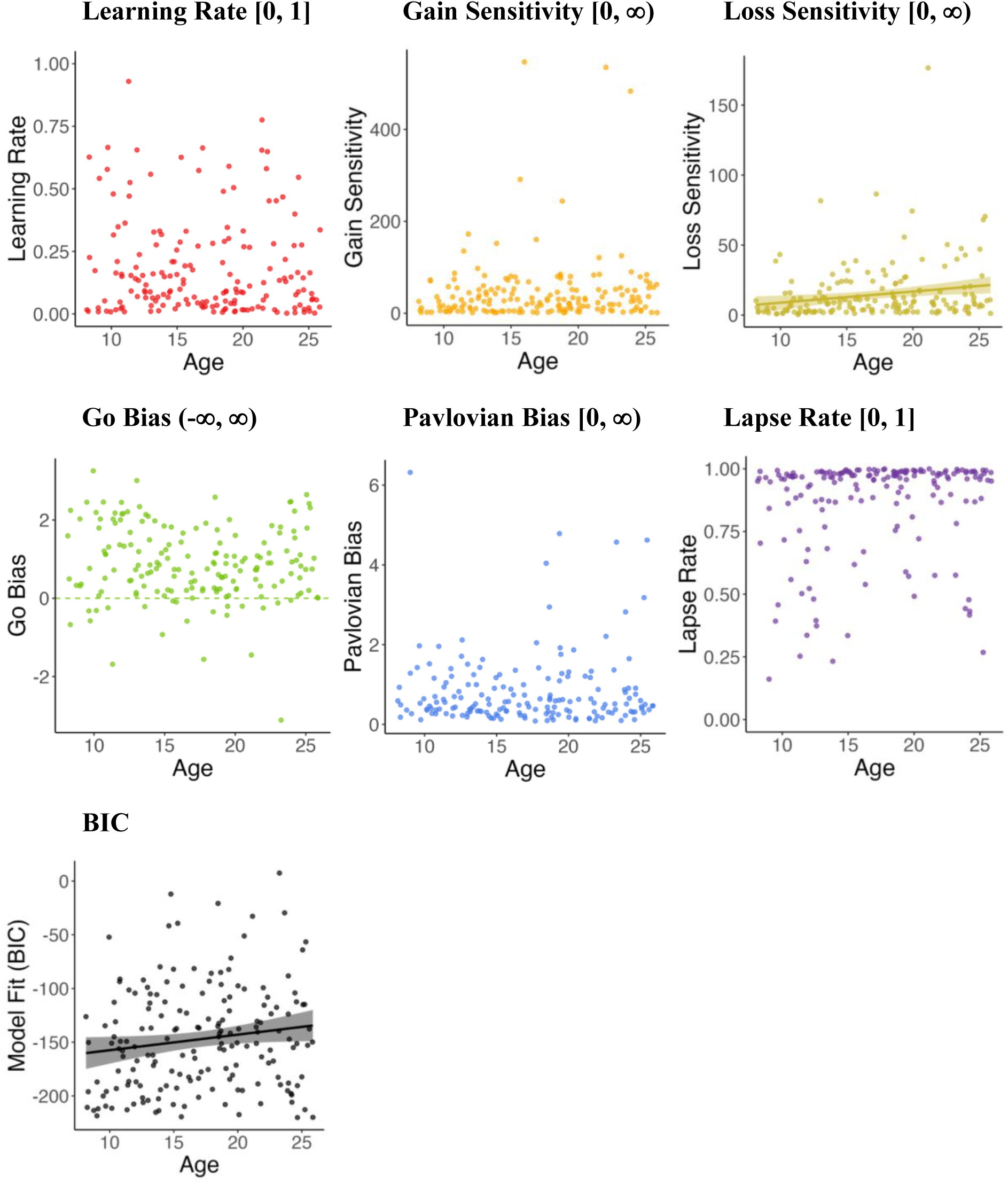
Parameter Associations with Age. We tested linear and quadratic associations with age for each parameter. The only parameter that significantly related to age is loss sensitivity, shown in the top right plot (Linear Age vs. Null Model: *F*(1,172) = 7.04, *p* < .01; *r* = .20, 95% CI: [.05, .34]), while the null model fit all other parameters best (Linear Age vs. Null Model *p*s > .05). Fit lines are shown for statistically significant effects. Points show individual-level parameter estimates, and shaded regions show 95% CIs.

### Memory Behavioral Analyses

**Figure S11.**
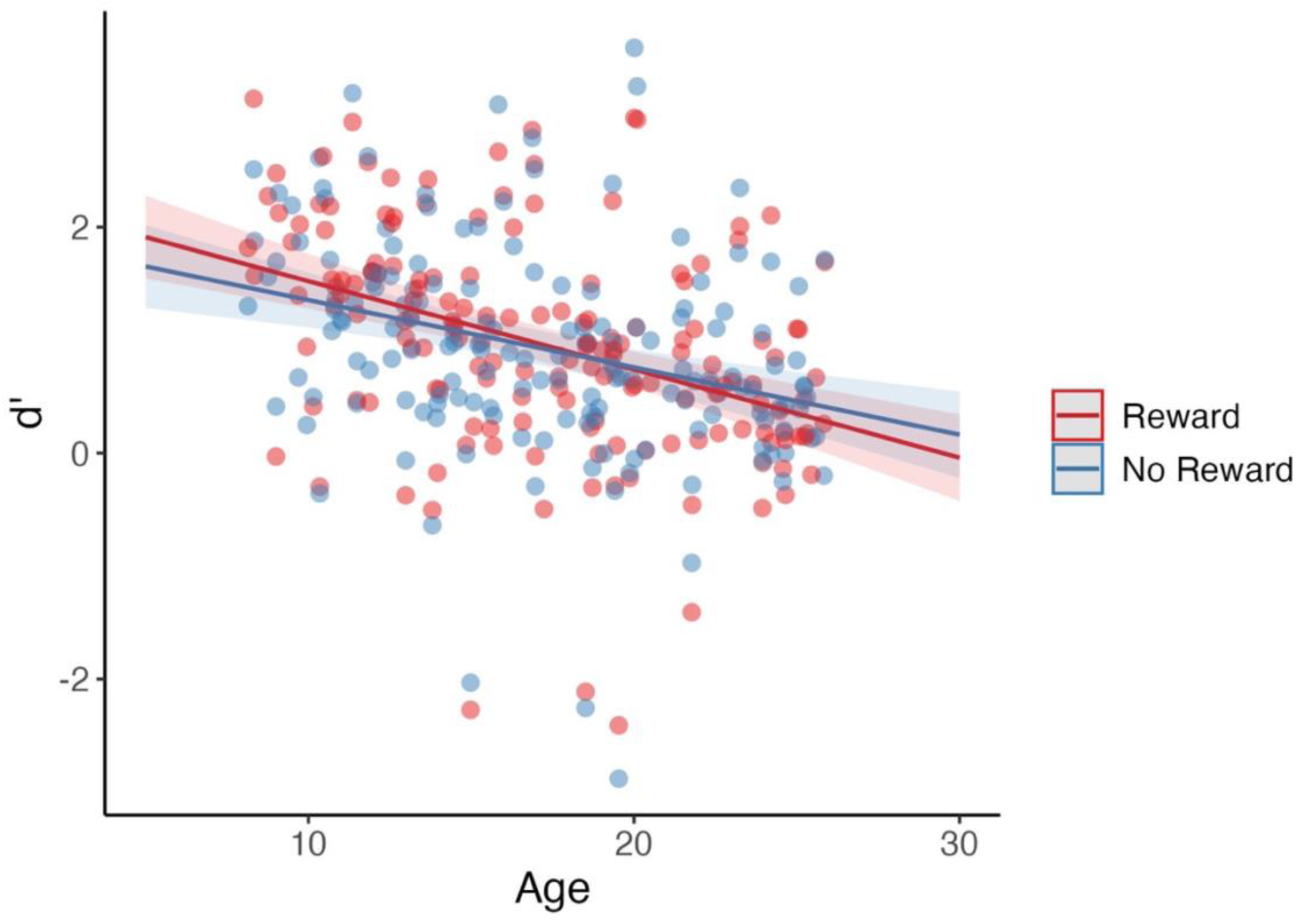
Effects Plot of Reward*Age Interaction Predicting d’. The lines represent the predicted values of the interaction of Reward*Age on d’. Memory accuracy of both types is worse in the older participants than in the younger. Younger participants remembered images associated with gaining rewards (red line) better than images associated with not gaining rewards (blue line). Older participants showed the opposite, remembering images associated with rewards relatively worse. The points show each participants’ sensitivity for each reward condition, and shaded regions show 95% confidence intervals.

**Figure S12.**
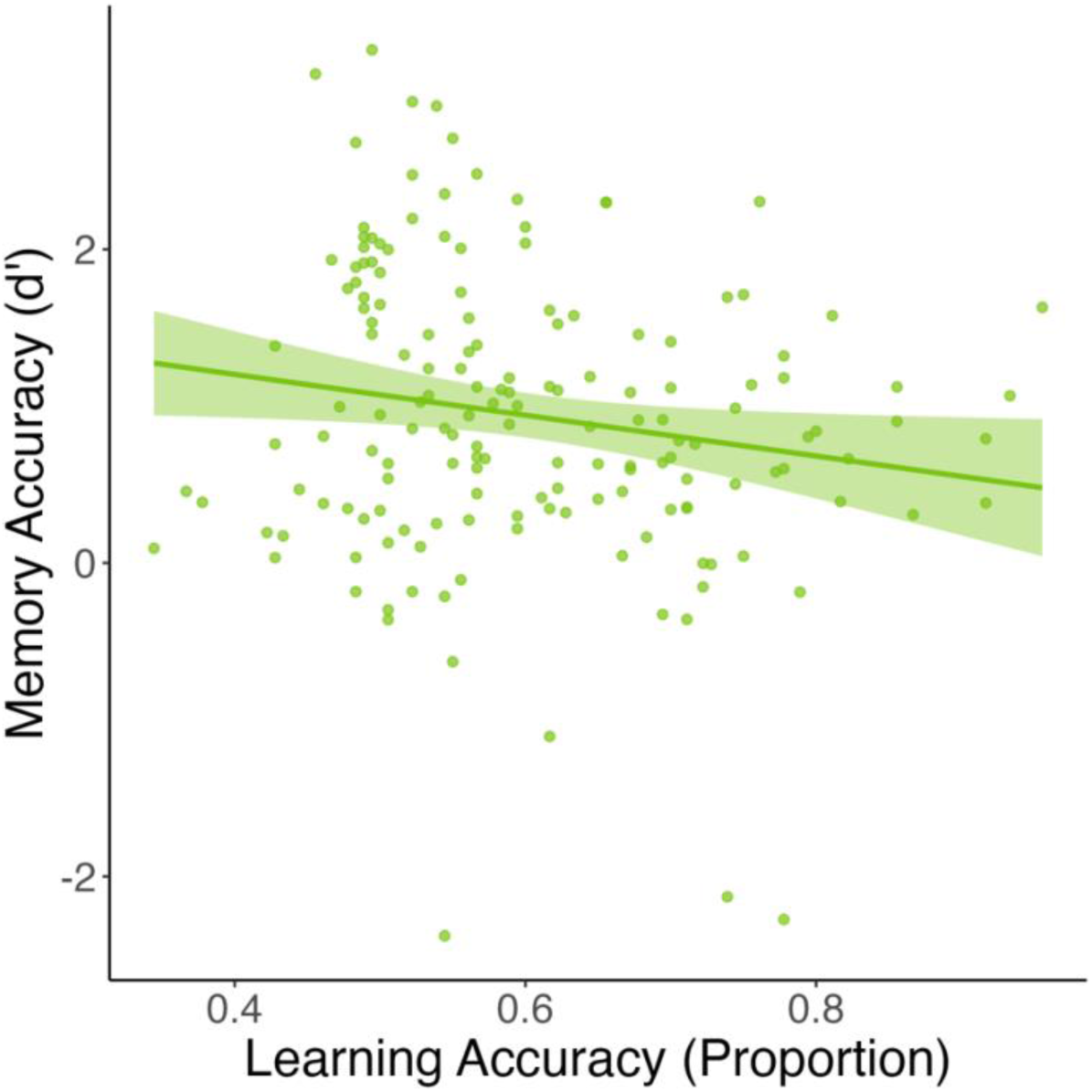
Association between Learning and Memory Accuracy. There is a weak, negative correlation between learning and memory accuracy (*r* = -.17, 95% CI = [−.32, -.02], *t*(160) = - 2.20, *p* = .03). The negative correlation suggests that better learning comes at the cost of incidental memory, which is generally consistent with prior work in adults (Wimmer et al., 2014).

**Table S5.**
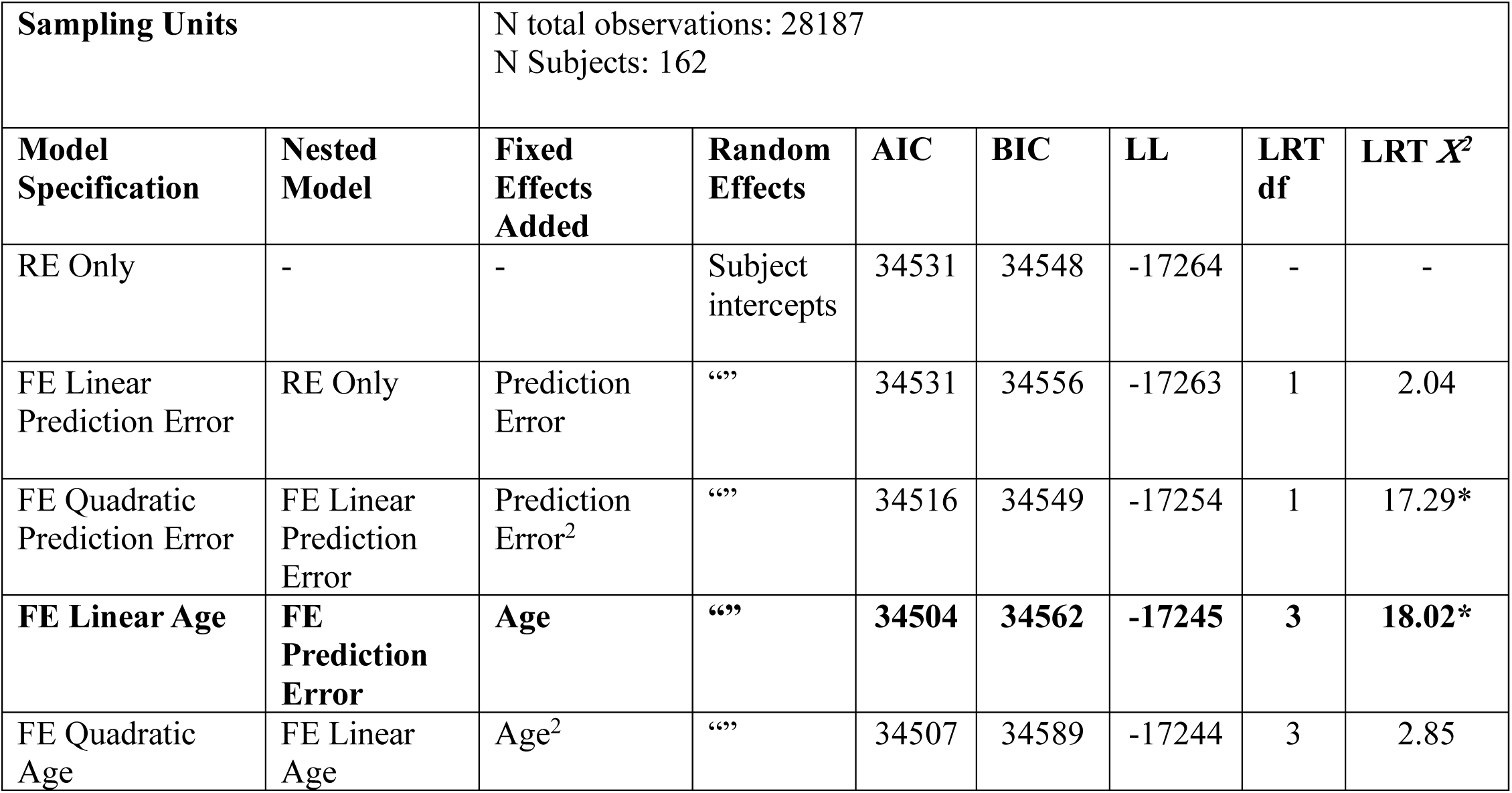
Model Comparison for Prediction Error-Related Memory Models. Likelihood ratio tests indicate that the best-fitting model includes linear and quadratic prediction error and linear age, the FE Linear Age model.

| Sampling Units |  | N total observations: 28187<br>N Subjects: 162 |  |  |  |  |  |  |
| --- | --- | --- | --- | --- | --- | --- | --- | --- |
| Model Specification | Nested Model | Fixed Effects Added | Random Effects | AIC | BIC | LL | LRT df | LRT $X^2$ |
| RE Only | - | - | Subject intercepts | 34531 | 34548 | -17264 | - | - |
| FE Linear Prediction Error | RE Only | Prediction Error | “” | 34531 | 34556 | -17263 | 1 | 2.04 |
| FE Quadratic Prediction Error | FE Linear Prediction Error | Prediction Error<sup>2</sup> | “” | 34516 | 34549 | -17254 | 1 | 17.29* |
| **FE Linear Age** | **FE Prediction Error** | **Age** | **“”** | **34504** | **34562** | **-17245** | **3** | **18.02\*** |
| FE Quadratic Age | FE Linear Age | Age<sup>2</sup> | “” | 34507 | 34589 | -17244 | 3 | 2.85 |

## REFERENCES

Adcock, R. A., Thangavel, A., Whitfield-Gabrieli, S., Knutson, B., & Gabrieli, J. D. E. (2006). Reward-Motivated Learning: Mesolimbic Activation Precedes Memory Formation. Neuron, 50(3), 507–517. 10.1016/j.neuron.2006.03.036

Aron, A. R., Robbins, T. W., & Poldrack, R. A. (2014). Inhibition and the right inferior frontal cortex: One decade on. Trends in Cognitive Sciences, 18(4), 177–185. 10.1016/j.tics.2013.12.003

Arsenault, J. T., Rima, S., Stemmann, H., & Vanduffel, W. (2014). Role of the Primate Ventral Tegmental Area in Reinforcement and Motivation. Current Biology, 24(12), 1347–1353. 10.1016/j.cub.2014.04.044

Averbeck, B., & O’Doherty, J. P. (2022). Reinforcement-learning in fronto-striatal circuits. Neuropsychopharmacology, 47(1), 147–162. 10.1038/s41386-021-01108-0

Barto, A. G., & Sutton, R. S. (1982). Simulation of anticipatory responses in classical conditioning by a neuron-like adaptive element. Behavioural Brain Research, 4(3), 221–235. 10.1016/0166-4328(82)90001-8

Ben-Shachar, M. S., Lüdecke, D., & Makowski, D. (2020). effectsize: Estimation of Effect Size Indices and Standardized Parameters. Journal of Open Source Software, 5(56), 2815. 10.21105/joss.02815

Betts, M. J., Richter, A., de Boer, L., Tegelbeckers, J., Perosa, V., Baumann, V., Chowdhury, R., Dolan, R. J., Seidenbecher, C., Schott, B. H., Düzel, E., Guitart-Masip, M., & Krauel, K. (2020). Learning in anticipation of reward and punishment: Perspectives across the human lifespan. Neurobiology of Aging, 96, 49–57. 10.1016/j.neurobiolaging.2020.08.011

Blass, E. M., Ganchrow, J. R., & Steiner, J. E. (1984). Classical conditioning in newborn humans 2–48 hours of age. Infant Behavior and Development, 7(2), 223–235. 10.1016/S0163-6383(84)80060-0

Bolenz, F., Reiter, A. M. F., & Eppinger, B. (2017). Developmental Changes in Learning: Computational Mechanisms and Social Influences. Frontiers in Psychology, 8. 10.3389/fpsyg.2017.02048

Bouton, M. E., & Bolles, R. C. (1980). Conditioned fear assessed by freezing and by the suppression of three different baselines. Animal Learning & Behavior, 8(3), 429–434. 10.3758/BF03199629

Braams, B. R., Van Duijvenvoorde, A. C. K., Peper, J. S., & Crone, E. A. (2015). Longitudinal Changes in Adolescent Risk-Taking: A Comprehensive Study of Neural Responses to Rewards, Pubertal Development, and Risk-Taking Behavior. The Journal of Neuroscience, 35(18), 7226–7238. 10.1523/JNEUROSCI.4764-14.2015

Brooks, M., E., Kristensen, K., Benthem, K., J., van Magnusson, A., Berg, C., W., Nielsen, A., Skaug, H., J., Mächler, M., & Bolker, B., M. (2017). glmmTMB Balances Speed and Flexibility Among Packages for Zero-inflated Generalized Linear Mixed Modeling. The R Journal, 9(2), 378. 10.32614/RJ-2017-066

Calderon, C. B., De Loof, E., Ergo, K., Snoeck, A., Boehler, C. N., & Verguts, T. (2021). Signed Reward Prediction Errors in the Ventral Striatum Drive Episodic Memory. The Journal of Neuroscience, 41(8), 1716–1726. 10.1523/JNEUROSCI.1785-20.2020

Casey, B. J., Cohen, A. O., & Galván, A. (2025). The beautiful adolescent brain: An evolutionary developmental perspective. Annals of the New York Academy of Sciences, 1546(1), 58–74. 10.1111/nyas.15314

Casey, B. J., Heller, A. S., Gee, D. G., & Cohen, A. O. (2019). Development of the emotional brain. Neuroscience Letters, 693, 29–34. 10.1016/j.neulet.2017.11.055

Cavanagh, J. F., Eisenberg, I., Guitart-Masip, M., Huys, Q., & Frank, M. J. (2013). Frontal Theta Overrides Pavlovian Learning Biases. The Journal of Neuroscience, 33(19), 8541–8548. 10.1523/JNEUROSCI.5754-12.2013

Chowdhury, R., Guitart-Masip, M., Lambert, C., Dolan, R. J., & Düzel, E. (2013). Structural integrity of the substantia nigra and subthalamic nucleus predicts flexibility of instrumental learning in older-age individuals. Neurobiology of Aging, 34(10), 2261–2270. 10.1016/j.neurobiolaging.2013.03.030

Cuevas, K., & Colombo, J. (2025). A quarter century of research on infant contingency learning: Current and future directions. Infant Behavior and Development, 80, 102068. 10.1016/j.infbeh.2025.102068

Davidow, J. Y., Foerde, K., Galván, A., & Shohamy, D. (2016). An Upside to Reward Sensitivity: The Hippocampus Supports Enhanced Reinforcement Learning in Adolescence. Neuron, 92(1), 93–99. 10.1016/j.neuron.2016.08.031

Davidow, J. Y., Insel, C., & Somerville, L. H. (2018). Adolescent Development of Value-Guided Goal Pursuit. Trends in Cognitive Sciences, 22(8), 725–736. 10.1016/j.tics.2018.05.003

Davidow, J. Y., Sheridan, M. A., Van Dijk, K. R. A., Santillana, R. M., Snyder, J., Vidal Bustamante, C. M., Rosen, B. R., & Somerville, L. H. (2019). Development of Prefrontal Cortical Connectivity and the Enduring Effect of Learned Value on Cognitive Control. Journal of Cognitive Neuroscience, 31(1), 64–77. 10.1162/jocn_a_01331

Dayan, P., Niv, Y., Seymour, B., & D. Daw, N. (2006). The misbehavior of value and the discipline of the will. Neural Networks, 19(8), 1153–1160. 10.1016/j.neunet.2006.03.002

Doremus-Fitzwater, T. L., Barreto, M., & Spear, L. P. (2012). Age-related differences in impulsivity among adolescent and adult Sprague-Dawley rats. Behavioral Neuroscience, 126(5), 735–741. 10.1037/a0029697

Dorfman, H. M., & Gershman, S. J. (2019). Controllability governs the balance between Pavlovian and instrumental action selection. Nature Communications, 10(1), 5826. 10.1038/s41467-019-13737-7

Eckstein, M. K., Master, S. L., Xia, L., Dahl, R. E., Wilbrecht, L., & Collins, A. G. (2022). The interpretation of computational model parameters depends on the context. eLife, 11, e75474. 10.7554/eLife.75474

Eldar, E., Hauser, T. U., Dayan, P., & Dolan, R. J. (2016). Striatal structure and function predict individual biases in learning to avoid pain. Proceedings of the National Academy of Sciences, 113(17), 4812–4817. 10.1073/pnas.1519829113

Ergo, K., Loof, E. D., & Verguts, T. (2020). Reward Prediction Error and Declarative Memory. Trends in Cognitive Sciences, 24(5), 388–397. 10.1016/j.tics.2020.02.009

Ferrari, A., & Comelli, M. (2016). A comparison of methods for the analysis of binomial clustered outcomes in behavioral research. Journal of Neuroscience Methods, 274, 131–140. 10.1016/j.jneumeth.2016.10.005

Fosco, W. D., Meisel, S. N., Weigard, A., White, C. N., & Colder, C. R. (2022). Computational modeling reveals strategic and developmental differences in the behavioral impact of reward across adolescence. Developmental Science, 25(2), e13159. 10.1111/desc.13159

Galván, A. (2010). Adolescent development of the reward system. Frontiers in Human Neuroscience, 4. 10.3389/neuro.09.006.2010

Galván, A., Hare, T. A., Parra, C. E., Penn, J., Voss, H., Glover, G., & Casey, B. J. (2006). Earlier Development of the Accumbens Relative to Orbitofrontal Cortex Might Underlie Risk-Taking Behavior in Adolescents. Journal of Neuroscience, 26(25), 6885–6892. 10.1523/JNEUROSCI.1062-06.2006

Gershman, S. J., Guitart-Masip, M., & Cavanagh, J. F. (2021). Neural signatures of arbitration between Pavlovian and instrumental action selection. PLOS Computational Biology, 17(2), e1008553. 10.1371/journal.pcbi.1008553

Guitart-Masip, M., Duzel, E., Dolan, R., & Dayan, P. (2014). Action versus valence in decision making. Trends in Cognitive Sciences, 18(4), 194–202. 10.1016/j.tics.2014.01.003

Guitart-Masip, M., Huys, Q. J. M., Fuentemilla, L., Dayan, P., Duzel, E., & Dolan, R. J. (2012). Go and no-go learning in reward and punishment: Interactions between affect and effect. NeuroImage, 62(1), 154–166. 10.1016/j.neuroimage.2012.04.024

Harden, K. P., & Tucker-Drob, E. M. (2011). Individual differences in the development of sensation seeking and impulsivity during adolescence: Further evidence for a dual systems model. Developmental Psychology, 47(3), 739–746. 10.1037/a0023279

Hart, G., Leung, B. K., & Balleine, B. W. (2014). Dorsal and ventral streams: The distinct role of striatal subregions in the acquisition and performance of goal-directed actions. Neurobiology of Learning and Memory, 108, 104–118. 10.1016/j.nlm.2013.11.003

Hartig, F. (2024). DHARMa: Residual Diagnostics for Hierarchical (Multi-Level / Mixed) Regression Models [Computer software]. https://CRAN.R-project.org/package=DHARMa

Hauser, T. U., Eldar, E., & Dolan, R. J. (2017). Separate mesocortical and mesolimbic pathways encode effort and reward learning signals. Proceedings of the National Academy of Sciences, 114(35). 10.1073/pnas.1705643114

Hauser, T. U., Iannaccone, R., Walitza, S., Brandeis, D., & Brem, S. (2015). Cognitive flexibility in adolescence: Neural and behavioral mechanisms of reward prediction error processing in adaptive decision making during development. NeuroImage, 104, 347–354. 10.1016/j.neuroimage.2014.09.018

Hauser, T. U., Will, G.-J., Dubois, M., & Dolan, R. J. (2019). Annual Research Review: Developmental computational psychiatry. Journal of Child Psychology and Psychiatry, 60(4), 412–426. 10.1111/jcpp.12964

Hershberger, W. A. (1986). An approach through the looking-glass. Animal Learning & Behavior, 14(4), 443–451. 10.3758/BF03200092

Huys, Q. J. M. (2018). Bayesian Approaches to Learning and Decision-Making. In Computational Psychiatry (pp. 247–271). Elsevier. 10.1016/B978-0-12-809825-7.00010-9

Huys, Q. J. M., Browning, M., Paulus, M. P., & Frank, M. J. (2021). Advances in the computational understanding of mental illness. Neuropsychopharmacology, 46(1), 3–19. 10.1038/s41386-020-0746-4

Huys, Q. J. M., Cools, R., Gölzer, M., Friedel, E., Heinz, A., Dolan, R. J., & Dayan, P. (2011). Disentangling the Roles of Approach, Activation and Valence in Instrumental and Pavlovian Responding. PLoS Computational Biology, 7(4), e1002028. 10.1371/journal.pcbi.1002028

Huys, Q. J. M., Gölzer, M., Friedel, E., Heinz, A., Cools, R., Dayan, P., & Dolan, R. J. (2016). The specificity of Pavlovian regulation is associated with recovery from depression. Psychological Medicine, 46(5), 1027–1035. 10.1017/S0033291715002597

Huys, Q. J. M., Maia, T. V., & Frank, M. J. (2016). Computational psychiatry as a bridge from neuroscience to clinical applications. Nature Neuroscience, 19(3), 404–413. 10.1038/nn.4238

Jang, A. I., Nassar, M. R., Dillon, D. G., & Frank, M. J. (2019). Positive reward prediction errors during decision-making strengthen memory encoding. Nature Human Behaviour, 3(7), 719–732. 10.1038/s41562-019-0597-3

Kass, R. E., & Raftery, A. E. (1995). Bayes Factors. Journal of the American Statistical Association, 90(430), 773–795. 10.2307/2291091

Lee, F. S., Heimer, H., Giedd, J. N., Lein, E. S., Šestan, N., Weinberger, D. R., & Casey, B. J. (2014). Adolescent mental health—Opportunity and obligation. Science, 346(6209), 547–549. 10.1126/science.1260497

Lüdecke, D. (2018). ggeffects: Tidy Data Frames of Marginal Effects from Regression Models. Journal of Open Source Software, 3(26), 772. 10.21105/joss.00772

Lüdecke, D. (2023). sjPlot: Data Visualization for Statistics in Social Science. R Package Version 2.8.15. https://CRAN.R-project.org/package=sjPlot

Lüdecke, D., Ben-Shachar, M., Patil, I., Waggoner, P., & Makowski, D. (2021). performance: An R Package for Assessment, Comparison and Testing of Statistical Models. Journal of Open Source Software, 6(60), 3139. 10.21105/joss.03139

Makowski, D. (2018). The Psycho Package: An Efficient and Publishing-Oriented Workflow for Psychological Science. Journal of Open Source Software, 3(22), 470.

Millner, A. J., Den Ouden, H. E. M., Gershman, S. J., Glenn, C. R., Kearns, J. C., Bornstein, A. M., Marx, B. P., Keane, T. M., & Nock, M. K. (2019). Suicidal thoughts and behaviors are associated with an increased decision-making bias for active responses to escape aversive states. Journal of Abnormal Psychology, 128(2), 106–118. 10.1037/abn0000395

Mkrtchian, A., Aylward, J., Dayan, P., Roiser, J. P., & Robinson, O. J. (2017). Modeling Avoidance in Mood and Anxiety Disorders Using Reinforcement Learning. Biological Psychiatry, 82(7), 532–539. 10.1016/j.biopsych.2017.01.017

Moutoussis, M., Bullmore, E. T., Goodyer, I. M., Fonagy, P., Jones, P. B., Dolan, R. J., & Dayan, P. (2018). Change, stability, and instability in the Pavlovian guidance of behaviour from adolescence to young adulthood. PLOS Computational Biology, 14(12), e1006679. 10.1371/journal.pcbi.1006679

Nussenbaum, K., & Hartley, C. A. (2019). Reinforcement learning across development: What insights can we draw from a decade of research? Developmental Cognitive Neuroscience, 40, 100733. 10.1016/j.dcn.2019.100733

O’Doherty, J., Dayan, P., Schultz, J., Deichmann, R., Friston, K., & Dolan, R. J. (2004). Dissociable Roles of Ventral and Dorsal Striatum in Instrumental Conditioning. Science, 304(5669), 452–454. 10.1126/science.1094285

Peirce, J., Gray, J. R., Simpson, S., MacAskill, M., Höchenberger, R., Sogo, H., Kastman, E., & Lindeløv, J. K. (2019). PsychoPy2: Experiments in behavior made easy. Behavior Research Methods, 51(1), 195–203. 10.3758/s13428-018-01193-y

Peters, S., & Crone, E. A. (2017). Increased striatal activity in adolescence benefits learning. Nature Communications, 8(1), 1983. 10.1038/s41467-017-02174-z

Pupillo, F., Ortiz-Tudela, J., Bruckner, R., & Shing, Y. L. (2023). The effect of prediction error on episodic memory encoding is modulated by the outcome of the predictions. Npj Science of Learning, 8(1), 18. 10.1038/s41539-023-00166-x

R Core Team. (2023). R: A Language and Environment for Statistical Computing. [Computer software]. R Foundation for Statistical Computing. https://www.R-project.org/

Raab, H. A., Goldway, N., Foord, C., & Hartley, C. A. (2024). Adolescents flexibly adapt action selection based on controllability inferences. Learning & Memory, 31(3), a053901. 10.1101/lm.053901.123

Raab, H. A., & Hartley, C. A. (2018). The Development of Goal-Directed Decision-Making. In R. Morris, A. Bornstein, & A. Shenhav (Eds.), Goal-Directed Decision Making: Computations and Neutral Circuits. Academic Press.

Raab, H. A., & Hartley, C. A. (2020). Adolescents exhibit reduced Pavlovian biases on instrumental learning. Scientific Reports, 10(1), 15770. 10.1038/s41598-020-72628-w

Rescorla, R. A., & Solomon, R. L. (1967). Two-process learning theory: Relationships between Pavlovian conditioning and instrumental learning. Psychological Review, 74(3), 151–182. 10.1037/h0024475

Rodman, A. M., Powers, K. E., Insel, C., Kastman, E. K., Kabotyanski, K. E., Stark, A. M., Worthington, S., & Somerville, L. H. (2021). How adolescents and adults translate motivational value to action: Age-related shifts in strategic physical effort exertion for monetary rewards. Journal of Experimental Psychology: General, 150(1), 103–113. 10.1037/xge0000769

Rosenbaum, G. M., Grassie, H. L., & Hartley, C. A. (2022). Valence biases in reinforcement learning shift across adolescence and modulate subsequent memory. eLife, 11, e64620. 10.7554/eLife.64620

Rouhani, N., & Niv, Y. (2021). Signed and unsigned reward prediction errors dynamically enhance learning and memory. eLife, 10, e61077. 10.7554/eLife.61077

Schreuders, E., Braams, B. R., Blankenstein, N. E., Peper, J. S., Güroğlu, B., & Crone, E. A. (2018). Contributions of Reward Sensitivity to Ventral Striatum Activity Across Adolescence and Early Adulthood. Child Development, 89(3), 797–810. 10.1111/cdev.13056

Schultz, W., Dayan, P., & Montague, P. R. (1997). A Neural Substrate of Prediction and Reward. Science, 275(5306), 1593–1599. 10.1126/science.275.5306.1593

Schwarz, G. (1978). Estimating the Dimension of a Model. The Annals of Statistics, 6(2), 461–464.

Seymour, B., Daw, N. D., Roiser, J. P., Dayan, P., & Dolan, R. (2012). Serotonin Selectively Modulates Reward Value in Human Decision-Making. The Journal of Neuroscience, 32(17), 5833–5842. 10.1523/JNEUROSCI.0053-12.2012

Somerville, L. H., Hare, T., & Casey, B. J. (2011). Frontostriatal Maturation Predicts Cognitive Control Failure to Appetitive Cues in Adolescents. Journal of Cognitive Neuroscience, 23(9), 2123–2134. 10.1162/jocn.2010.21572

Somerville, L. H., Jones, R. M., Ruberry, E. J., Dyke, J. P., Glover, G., & Casey, B. J. (2013). The Medial Prefrontal Cortex and the Emergence of Self-Conscious Emotion in Adolescence. Psychological Science, 24(8), 1554–1562. 10.1177/0956797613475633

Stanislaw, H., & Todorov, N. (1999). Calculation of signal detection theory measures. Behavior Research Methods, Instruments, & Computers, 31(1), 137–149. 10.3758/BF03207704

Steinberg, L., Albert, D., Cauffman, E., Banich, M., Graham, S., & Woolard, J. (2008). Age Differences in Sensation Seeking and Impulsivity as Indexed by Behavior and Self-Report: Evidence for a Dual Systems Model. Developmental Psychology, 44, 1764–1778. 10.1037/a0012955

Störmer, V., Eppinger, B., & Li, S.-C. (2014). Reward speeds up and increases consistency of visual selective attention: A lifespan comparison. Cognitive, Affective, & Behavioral Neuroscience, 14(2), 659–671. 10.3758/s13415-014-0273-z

Sutton, R. S., & Barto, A. G. (1981). Toward a modern theory of adaptive networks: Expectation and prediction. Psychological Review, 88(2), 135–170. 10.1037/0033-295X.88.2.135

Sutton, R. S., & Barto, A. G. (1998). Reinforcement learning: An introduction. MIT Press.

Tremblay, L., Hollerman, J. R., & Schultz, W. (1998). Modifications of Reward Expectation-Related Neuronal Activity During Learning in Primate Striatum. Journal of Neurophysiology, 80(2), 964–977. 10.1152/jn.1998.80.2.964

Tummeltshammer, K., Feldman, E. C. H., & Amso, D. (2019). Using pupil dilation, eye-blink rate, and the value of mother to investigate reward learning mechanisms in infancy. Developmental Cognitive Neuroscience, 36, 100608. 10.1016/j.dcn.2018.12.006

Uhlhaas, P. J., Davey, C. G., Mehta, U. M., Shah, J., Torous, J., Allen, N. B., Avenevoli, S., Bella-Awusah, T., Chanen, A., Chen, E. Y. H., Correll, C. U., Do, K. Q., Fisher, H. L., Frangou, S., Hickie, I. B., Keshavan, M. S., Konrad, K., Lee, F. S., Liu, C. H., … Wood, S. J. (2023). Towards a youth mental health paradigm: A perspective and roadmap. Molecular Psychiatry, 28(8), 3171–3181. 10.1038/s41380-023-02202-z

van den Bos, W., Cohen, M. X., Kahnt, T., & Crone, E. A. (2012). Striatum–Medial Prefrontal Cortex Connectivity Predicts Developmental Changes in Reinforcement Learning. Cerebral Cortex, 22(6), 1247–1255. 10.1093/cercor/bhr198

van Duijvenvoorde, A. C. K., Westhoff, B., de Vos, F., Wierenga, L. M., & Crone, E. A. (2019). A three-wave longitudinal study of subcortical–cortical resting-state connectivity in adolescence: Testing age- and puberty-related changes. Human Brain Mapping, 40(13), 3769–3783. 10.1002/hbm.24630

Van Leijenhorst, L., Zanolie, K., Van Meel, C. S., Westenberg, P. M., Rombouts, S. A. R. B., & Crone, E. A. (2010). What Motivates the Adolescent? Brain Regions Mediating Reward Sensitivity across Adolescence. Cerebral Cortex, 20(1), 61–69. 10.1093/cercor/bhp078

Wassum, K. M., Ostlund, S. B., Balleine, B. W., & Maidment, N. T. (2011). Differential dependence of Pavlovian incentive motivation and instrumental incentive learning processes on dopamine signaling. Learning & Memory, 18(7), 475–483. 10.1101/lm.2229311

Wechsler, D. (2011). Wechsler Abbreviated Scale of Intelligence—Second Edition [Dataset]. 10.1037/t15171-000

Wickham, H. (2016). ggplot2: Elegant Graphics for Data Analysis (Vol. 67). Springer-Verlag New York. https://ggplot2.tidyverse.org

Wickham, H., Averick, M., Bryan, J., Chang, W., McGowan, L., François, R., Grolemund, G., Hayes, A., Henry, L., Hester, J., Kuhn, M., Pedersen, T., Miller, E., Bache, S., Müller, K., Ooms, J., Robinson, D., Seidel, D., Spinu, V., … Yutani, H. (2019). Welcome to the Tidyverse. Journal of Open Source Software, 4(43), 1686. 10.21105/joss.01686

Wilbrecht, L., & Davidow, J. Y. (2024). Goal-directed learning in adolescence: Neurocognitive development and contextual influences. Nature Reviews Neuroscience, 25(3), 176–194. 10.1038/s41583-023-00783-w

Wiltgen, B. J., Law, M., Ostlund, S., Mayford, M., & Balleine, B. W. (2007). The influence of Pavlovian cues on instrumental performance is mediated by CaMKII activity in the striatum. European Journal of Neuroscience, 25(8), 2491–2497. 10.1111/j.1460-9568.2007.05487.x

Wimmer, G. E., Braun, E. K., Daw, N. D., & Shohamy, D. (2014). Episodic Memory Encoding Interferes with Reward Learning and Decreases Striatal Prediction Errors. The Journal of Neuroscience, 34(45), 14901–14912. 10.1523/JNEUROSCI.0204-14.2014

Yin, H. H., Ostlund, S. B., Knowlton, B. J., & Balleine, B. W. (2005). The role of the dorsomedial striatum in instrumental conditioning. European Journal of Neuroscience, 22(2), 513–523. 10.1111/j.1460-9568.2005.04218.x

